# A global survey of small RNA interactors identifies KhpA and KhpB as major RNA-binding proteins in *Fusobacterium nucleatum*

**DOI:** 10.1101/2023.10.30.564711

**Authors:** Yan Zhu, Falk Ponath, Valentina Cosi, Jörg Vogel

## Abstract

The common oral microbe *Fusobacterium nucleatum* has recently drawn attention after it was found to colonize tumors throughout the human body. Fusobacteria are also interesting study systems for bacterial RNA biology as these early-branching species encode many small noncoding RNAs (sRNAs) but lack homologs of the common RNA-binding proteins (RBPs) CsrA, Hfq and ProQ. To search for alternate sRNA-associated RBPs in *F. nucleatum*, we performed a systematic mass spectrometry analysis of proteins that co-purified with 19 different sRNAs. This approach revealed strong enrichment of the KH domain proteins KhpA and KhpB with nearly all tested sRNAs, including the σ^E^-dependent sRNA FoxI, a regulator of several envelope proteins. KhpA/B act as a dimer to bind sRNAs with low micromolar affinity and influence the stability of several of their target transcripts. Transcriptome studies combined with biochemical and genetic analyses suggest that KhpA/B have several physiological functions, including being required for ethanolamine utilization. Our RBP search and the discovery of KhpA and KhpB as major RBPs in *F. nucleatum* are important first steps in identifying key players of post-transcriptional control at the root of the bacterial phylogenetic tree.

## INTRODUCTION

The anaerobic Gram-negative bacterium *Fusobacterium nucleatum* is a major constituent of the human oral microbiota (1). Long viewed primarily as one of several oral commensals, *F. nucleatum* has recently received attention because it was reported to spread from its primary niche to extra-oral sites. *F. nucleatum* has been shown to associate with several types of human cancers, including breast and colorectal cancer (Kostic et al., 2012; Parhi et al., 2020) and its presence appears to contribute to tumor progression (1,2) This raises the intriguing question how *F. nucleatum*, which is adapted to the oral cavity, can migrate to and survive in these secondary niches. It is fair to assume that this oral bacterium possesses gene regulation mechanisms that allow it to thrive in these very different environments, but to date, very little is known about these mechanisms. The last two years have seen the first reports of factors involved in transcriptional control. These include the two-components systems (TCSs) CarRS and ModRS, which regulate lysine metabolism and interspecies coaggregation (3) or defense against oxidative stress (4), respectively. Additionally, a homolog of the extracytoplasmic function (ECF) sigma factor σ^E^ has been reported, which in *F. nucleatum* seems to respond to oxygen stress to trigger a global envelope stress response (5).

In addition to the medical interest, *F. nucleatum* and fusobacteria in general are relevant study systems for RNA biology, as they constitute an early-branching phylum (the Fusobacteriota) that is phylogenetically remote from conventional model bacteria, including Proteobacteria, Bacteroidota and Firmicutes (6). Thus, studying small noncoding RNAs (sRNAs) and RNA-binding proteins (RBPs), two major classes of factors that mediate post-transcriptional control, is a promising strategy to obtain insights into the evolution of RNA-mediated gene regulation in bacteria. Using differential RNA-seq, we have previously identified 24 sRNAs in *F. nucleatum* (7). Our characterization of the sRNA FoxI showed that it is induced by σ^E^ under oxygen stress. It base-pairs to multiple target mRNAs to repress several envelope proteins, such as the inner membrane protein complex *mglBAC* and the outer-membrane protein *fomA* (5,7). FoxI is present in all *F. nucleatum* strains as well as in the two related strains *F. periodonticum* and *F. hwasookii* and harbors a conserved seed region for mRNA recognition (7). σ^E^-dependent sRNAs from *Escherichia coli* and *Salmonella enterica* rely on protein-assisted base-pairing to mRNA targets to modulate gene regulation (8,9), however, it has remained unclear if there are RBPs that act in concert with FoxI and other fusobacterial sRNAs.

To date, three proteins have been established as major sRNA-associated RBPs in Gram-negative bacteria: CsrA (a.k.a. RsmA) and the two RNA chaperones Hfq and ProQ. Collectively, these three RBPs mediate post-transcriptional control and broadly impact bacterial physiology and/or virulence in *E. coli*, *Pseudomonas aeruginosa*, *S. enterica, Vibrio cholerae,* and many other Gram-negative species (10–16). CsrA is widely conserved and was initially discovered as a carbon storage regulator (17). It binds to GGA motifs around the Shine-Dalgarno (SD) sequence of mRNAs to activate or repress translation (18,19) and is itself regulated by decoy sRNAs (20). Hfq is an RNA chaperone in Gram-negative bacteria and contributes to post-transcriptional regulation mediated by *trans*-encoded sRNAs (21). It recognizes A-rich sequences in the 5’ UTRs of mRNAs and U-rich sequences at the 3’ end of sRNAs to promote base-pairing between sRNA-mRNA pairs (18,22,23). ProQ was recently identified as an additional major RNA chaperone and binds to the 3’ UTRs of mRNAs and the 3’ end of sRNAs via a structure-driven mechanism. ProQ has also been shown to promote base-pairing between sRNAs and mRNAs (24–26). Importantly, fusobacteria are predicted to encode no homologs of CsrA, Hfq, or ProQ, raising the possibility that reported sRNAs in this phylum interact with yet unknow RBPs.

To identify RBPs in bacteria, different strategies have been developed (27–29). One is the tagging of RNAs with an aptamer or other sequence tags, so that they can be used as molecular baits to co-purify associated proteins from cellular lysates (30). For example, the 14-mer capture tag is specifically recognized by a complementary RNA adaptor, which is biotinylated and attached to streptavidin beads. 14-mer sequence tagged-RNAs are thus immobilized to capture RNA-interactors, which can be identified by mass spectrometry (MS) (Figure 1A) (31). Previously, we have successfully used this capture tag in *Streptococcus pneumoniae* to identify Cbf1 as a sRNA-associated RBP (32). Using *in vitro* synthesized tagged RNAs allows the application of this approach to non-model organisms that lack genetically tractable tools and due to experimental ease and low cost of RNA tagging, it is also feasible to screen multiple RNA baits in parallel (33).

**Figure 1.**
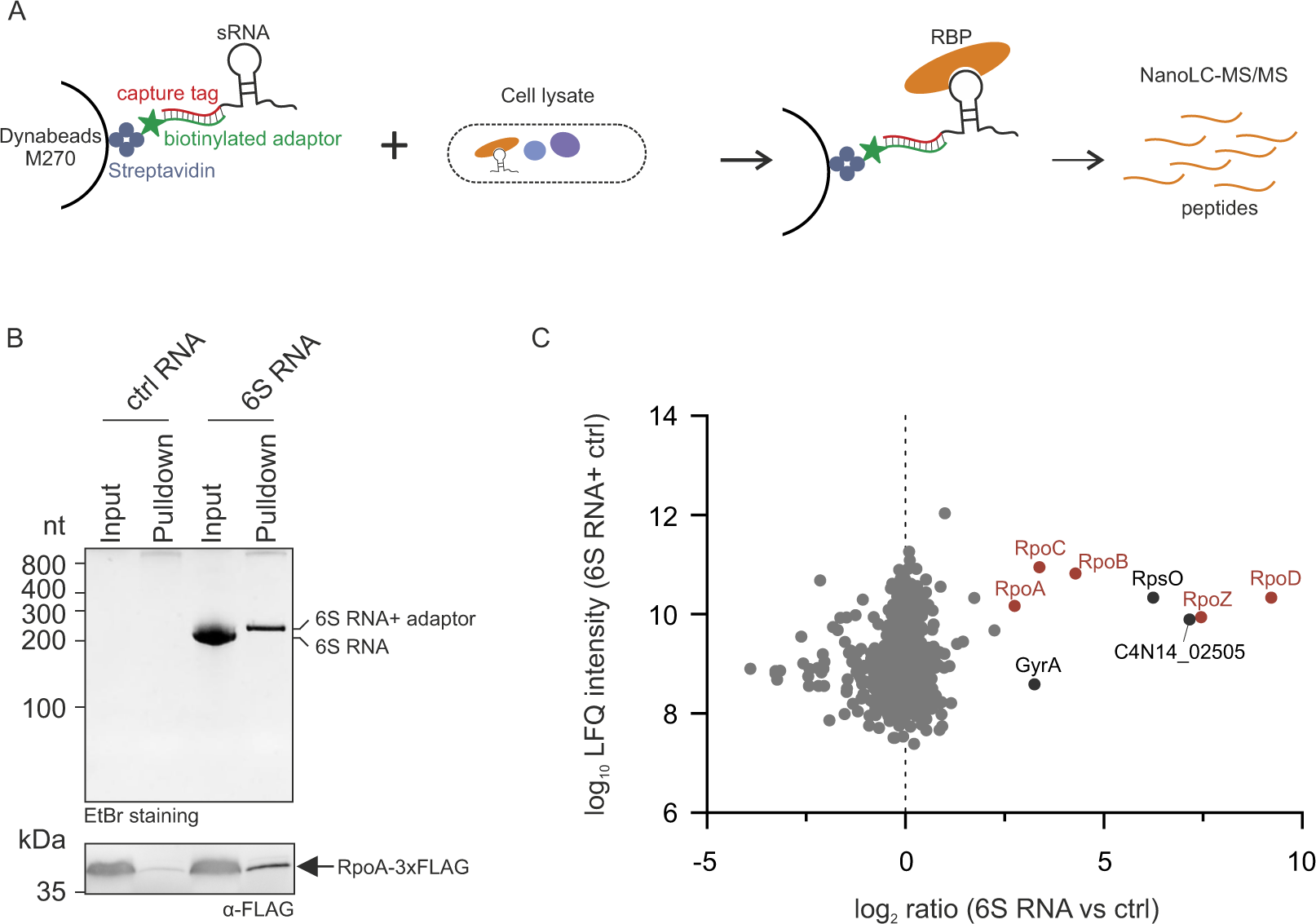
An RNA-centric approach to identify RNA-protein interactions in *F. nucleatum*. (**A**) Schematic representation of the 14 mer capture tagged-sRNA affinity purification strategy. (**B**) Gel images of the sRNA affinity purification performed with tagged 6S RNA incubated with a lysate of a *F. nucleatum* strain that expresses 3xFLAG-tagged RpoA. Upper panel: pulldown samples were separated on 8% acrylamide gel containing 8 M urea with 1 µg *in vitro* transcribed 6S RNA as input control. The gel was stained with Ethidium bromide. Lower panel: Western blot analysis of pulldown samples with anti-FLAG antibody. (**C**) Mass spectrometric analysis of the sRNA affinity purification using tagged-6S RNA as bait, with a F. nucleatum WT lysate. Log_10_ LFQ intensities of the 6S RNA and the control RNA are plotted against the log_2_ ratio between the 6S RNA versus the control RNA. RNA polymerase components (red) and three highly enriched proteins are labeled.

In the present work, we have tagged 19 *F. nucleatum* sRNAs and used them as baits in sRNA affinity purifications to identify a multitude of interesting RBP candidates. This included the KH domain proteins KhpA and KhpB, which act as broad sRNA-associated RBPs, echoing recent findings in phylogenetically distant Gram-positive bacteria (34–36). Our study further shows that in *F. nucleatum* KhpA and KhpB are global regulators involved in various physiological processes, including ethanolamine utilization. Together, these data lay the foundation for future studies of post-transcriptional regulation in *F. nucleatum*.

## MATERIAL AND METHODS

### Bacterial strains, plasmids and genetic manipulations

The bacterial strains and plasmids used in this study are listed in Supplementary Table S4. *Fusobacterium nucleatum* subspecies *nucleatum* ATCC 23726 and derivative strains are normally cultured in Columbia broth medium (BD) or on brain-heart infusion (BHI) 2% agar plates supplemented with 1% (w: v) yeast extract, 1% (w: v) glucose, 5 µg/ml of hemin and 1% (v: v) fetal bovine serum at 37 °C in anaerobic chamber (N_2_:H_2_:CO_2_=80%:10%:10%, Coy Laboratory Products). For the condition of nutrient limitation, the CYG medium (Casein peptone 10 mg/ml, HEPES 16.8 mM, Hemin 0.5 mg/ml, NaCl 5 mg/ml, Yeast 5 mg/ml, pH was adjusted with NaOH to 7.0) is used. For growth curve assay, 24 h pre-cultures were inoculated in a 96-well plate to an initial OD_600_ of 0.01. Measurements were performed with a plate reader (Biotek) in the anaerobic chamber.

Deletion strains and epitope tagged protein strain were generated with the pVoPo-04 system for markerless genomic deletion as previously described (5). Similarly, complementary strains were constructed with pVoPo-5 system. Briefly, approx. 1.2 kb of homology regions to the chromosomal sequence flanking the up- and down-stream of deletion or insertion sites were inserted into the pVoPo-04 backbone or pVoPo-05 backbone by Gibson assembly (NEBuilder HiFi DNA Assembly Master Mix, New England BioLabs). 5-10 μg of the constructed plasmids were dialyzed and electroporated into *F. nucelatum*. *mazF* induction with anhydrotetracycline (ATc) and double cross-over homologous recombination screen were performed as described (5). When used the pVoPo-04 and pVoPo-05 system in *F. nucleatum*, thiamphenicol was added with the concentration of 5 µg/ml in liquid medium and 2.5 µg/ml in agar plates. Oligonucleotides used for the construction of pVoPo-04- and pVoPo-05-based plasmids, PCR checking of deletion and insertion are listed in the Supplementary Table S4.

### In vitro synthesis of sRNA baits, control RNA and adaptor for sRNA affinity purification

sRNA baits used in sRNA affinity purification were *in vitro* transcribed using MEGAscript T7 Transcription kit (Thermo Fisher Scientific). Firstly, the desired DNA sequences were amplified from ATCC 23726 genomic DNA via specific primers (listed in Supplementary Table S4). An extension containing T7 promoter followed by the 14 nt tag was added in the 5’ PCR primer (5’- GTTTTTTTTAATACGACTCACTATAGG GAGACCTAGCCT -3’). 200 ng of the PCR-product was then used for the *in vitro* transcription reaction. RNA products were separated by 8% acrylamide gels containing 8 M urea. RNA bands were cut out and eluted in RNA elution buffer (0.1 M sodium acetate, 0.1% SDS, 10 mM EDTA) overnight at 4°C with rotation. Eluted RNAs were further extracted via Phenol: Chloroform: Isoamylalcohol (P: C: I, 25: 24: 1, pH 4.5, Roth) and precipitated with ethanol. The concentration of RNAs were measured with Nanodrop. The integrity of RNAs were checked by loading 1 µg on 8% acrylamide gels containing 8 M urea. The adaptor oligonucleotide (5’-AGGCUAGGUCUCCC-3’) which are complementary to the tag added in RNAs, was synthesized with all bases are 2’ -O-methyl modified and 3’-end is biotin modified (eurofins). The control oligo RNA harboring the tag (GGGAGACCUAGCCU) was synthesized as well (eurofins).

### sRNA affinity purification

For sRNA affinity purification, 100 μl of magnetic streptavidin beads (Dynabeads Streptavidin M- 270) were pre-washed 3 times with wash buffer (20 mM Tris-HCl pH 8.0, 150 mM KCl, 1 mM MgCl_2_, 5 % Glycerol, 0.01 % Tween-20). 4 μg of the adaptor oligonucleotide was added to the pre-washed beads and incubated for 1 h at 4 °C with rotation. After washing out the uncoupled adaptors with wash buffer, transfer 50 μl of the beads to a new tube containing 10 μg of bait-RNA or control RNA, another 50 μl to a *F. nucleatum* lysate for pre-clearing the lysate. To prepare the lysate, 200 ml of cultures from mid-exponential growth phase (OD_600_=0.5) were harvested and the pellet were resuspended in lysis buffer (20 mM Tris-HCl pH 8.0, 150 mM KCl, 1 mM MgCl_2_, 5% Glycerol, 0.01% Tween-20) and lysed by Retsch MM200 with 0.1 mm glass beads at 30 Hz for 10 min. Centrifuge the lysate and collect the supernatant to adaptor-coupled beads and rotate for 3.5 h at 4 °C. The RNA-adaptor-coupled beads were then incubated with the pre-cleared *F. nucleatum* lysate for 2 h at 4 °C. After washing the beads to get rid of unspecific binders, the elution fractions were collected for further analysis.

When the pulldowns were performed with 6S RNA or FoxI and FunR2 with western blot analysis for checking proteins recovery, the beads were split into two portions before last step of washing. After discard the wash buffer, one portion of beads was resuspended in 35 μl GLII buffer and boiled for 5 min at 95 °C and then loaded on 8% acrylamide gels containing 8 M urea with Ethidium bromide staining; 1 μg of 6S RNA was loaded as input control. Another portion was resuspended in 35 μl 1x LDS buffer (Thermo Fisher Scientific) containing 50 mM DTT and boiled for 5 min at 95 °C and loaded on 15 % SDS-PAGE gel for further western blot analysis. When the pulldowns were performed for subsequent MS analysis, after the last step of washing the beads were resuspended in 35 μl 1x LDS buffer (Thermo Fisher Scientific) containing 50 mM DTT and boiled for 5 min at 95 °C. The beads were quickly spun down and the supernatant was transferred into a fresh tube with 4.73 µL 1 M iodoacetamide for alkylation of the protein samples. After incubated for 20 min at room temperature, the samples were subjected to the Nano LC-MS/MS analysis.

### Nano LC-MS/MS analysis

Nano LC-MS/MS analysis for elute protein samples from sRNA affinity purification was performed as described previously (32) at Rudolf-Virchow-Centre Würzburg for Integrative and Translation Bioimaging. Briefly, the elute samples were destained in 100 mM ammonium bicarbonate containing 30% acetonitrile and shrunk in 100% acetonitrile before trypsin digestion (0.1 μg in 100 mM ammonium bicarbonate, overnight at 37 °C) and were further dissolved in 5% formic acid for subsequent nano-liquid chromatography–tandem MS (LC-MS/MS) analysis. For nanoLC–MS/MS analysis, peptides separation was performed by C-18 reversed phase chromatography (capillary columns, PicoFrit, 30 cm × 150 µm ID, New Objective packed with ReproSil-Pur 120 C18-AQ, 1.9 µm, Dr. Maisch), which is connected with an Orbitrap Fusion ETD (Thermo Scientific) equipped with a PicoView Ion Source (New Objective) and a nEASY- LC1000 liquid chromatography system (Thermo Scientific). Separation was performed using a 140-min linear gradient (3-40% acetonitrile, 0.1% formic acid) at a flow rate of 500 nL/min. Both MS and MS/MS analyses were conducted using the Orbitrap analyzer with a resolution of 60, 000 for MS scans and 15, 000 for MS/MS scans and the higher-energy collisional dissociation fragmentation was set to 35% normalized collision energy. A fixed cycle time of 3 sec was used for a Top Speed data-dependent MS/MS method. Dynamic exclusion with one repeat count per 60 sec was applied. Single charged precursors were excluded. The minimum threshold for precursors selection was set to 50,000. Internal calibration with Easy-IC was used to improve mass:charge ratio assignment.

Subsequent analysis of the data was conducted using MaxQuant (v.1.5.7.4) with integrated Andromeda comparing it against the Uniprot database for *F. nucleatum* subsp. *nucleatum* ATCC 23726.

### Protein purification

For recombinant protein purification, the coding region of full length KhpA or KhpB-ΔN (aa 83- 245) with a start codon ATG was cloned into the *E. coli* recombinant protein expressing vector pET28a with a 6xHis tag and a 3C protease cleavage site added in the N-terminus. The oligonucleotides used for the constructed are listed in Supplementary Table S4.

The purifications of KhpA and KhpB-ΔN were performed by the Recombinant Protein Expression core unit at the Rudolf Virchow Centre at the University of Würzburg. Briefly, the protein expression plasmids were transformed into *E. coli* BL21 (DE3) codonplus. The cells were grown to an OD_600_ of 0.5 at 37 °C and induced with 0.5 mM IPTG at 18 °C for overnight. Cell cultures were collected and cell pellets were resuspended in lysis buffer (50 mM Tris-HCl pH 8.0, 300 mM KCl, 10 % sucrose, 1 mM MgCl_2_, 10 mM imidazole, 1 mM TCEP) and lysed by sonication. The soluble lysate was incubated with Ni-NTA agarose and then subjected to column chromatography. Following eluted from the column with elution buffer (50 mM Tris-HCl pH 8.0, 300 mM KCl, 10 % sucrose, 1 mM MgCl_2_, 250 mM imidazole, 1 mM TCEP), the recombinant KhpA or KhpB protein was further incubated with 3C protease for His tag cleavage. At the end, the KhpA and KhpB-ΔN were secondly purified by cation-exchange chromatography (HiTrap SP 5 ml, GE Healthcare Life Science) followed by SDS-PAGE. The purified KhpA and KhpB-ΔN were aliquoted and stored at -80°C.

### Preparation and labeling of RNA

sRNAs used for the EMSAs were *in vitro* transcribed with the MEGAscript T7 transcription kit (Thermo Scientific) as for sRNAs baits used in the affinity purification. Then, 50 pmol of *in vitro* transcribed RNA was dephosphorylated with quick-CIP (New England Biolands) at 37 °C for 15 min and subsequently extracted with P: C: I. For radioactivity labeling the dephosphorylated RNAs, 20 pmol of CIP-treated RNA was incubated with 20 μCi 32_P_-γ-ATP and T4 polynucleotide kinase (PNK, Thermo Scientific) in a 20 μl reaction system at 37°C for 1 h. Labeled RNAs were then purified with Miscrospin G-50 columns (GE Healthcare) and stored at -20°C for EMSA assay later.

### EMSA

For EMSA assay, labeled RNAs and yeast tRNA (Ambion, in 1: 10 dilution) was denatured by heating for 1 min at 95°C and cooling on ice for 1min. 4 nM of each labeled RNA and 1 μg of yeast tRNA were incubated with increasing concentration (0, 0.16, 0.32, 0.63, 1.25, 2.5, 5 μM) of KhpA, KhpB-ΔN or both the proteins(1: 1 ratio) in binding buffer (25 mM Tris-HCl pH 7.4, 150 mM NaCl, 1 mM MgCl_2_) at 37°C for 1h. The reactions were stopped by adding 5x RNA native loading buffer and loading on a pre-cooled native 6% polyacrylamide gel and separated by running the gel at 4 °C in 0.5% TBE at constant current of 40 mA for 4h. After running, the gel was vacuum dried, and signals were detected on a Typhoon FLA 700 phosphoimager (GE Healthcare life science).

### Total RNA extraction and northern blot

*F. nucleatum* strains used were cultured to the desired conditions and a biomass of 6 OD_600_ of culture were collected and added with 2% stop mix (95% ethanol, 5% phenol) and snap-frozen in liquid nitrogen. The total RNAs were extracted with hot phenol method followed by DNase treatment as described previously (7). For northern blot, 5 µg of total RNAs were loaded on 6% acrylamide gels containing 8 M urea, electrophoretic separated in Tris–borate–EDTA (TBE) buffer. RNAs were transferred to Hybond N+ (GE Healthcare) filters. 5’ radiolabeled oligos were used as probes detecting transcripts of interest or 5S RNA which served as loading control. For oligonucleotide labeling, 10 pmol of the oligonucleotides used as probes were 5’-labeled with 10 µCi of 32^P^-*γ*-ATP by PNK (T4 polynucleotide kinase, Thermo Fisher Scientific) at 37 °C. Labeled oligonucleotides were further purified with Microspin G-25 columns (GE Healthcare) to remove unincorporated 32^P^-*γ*-ATP. Radioactive signal was imaged with the Typhoon FLA 7000 (GE Healthcare). Oligonucleotides used as probes in rifampicin assay and RNA immunoprecipitation are listed in Supplementary Table S4.

### total RNA-seq and data analysis

For total RNA-seq, the cDNA libraries preparation and sequencing were performed by the CoreUnit Sysmed at the University of Würzburg. Briefly, the rRNA depleted RNA was fragmented via ultrasound, 1 pulse of 30 seconds at 4 °C. An adapter was ligated to the 3’ end of fragmented RNA. The first-strand cDNA was synthesized using the M-MLV reverse transcriptase in conjunction with the introduced 3’-adapter serving as a primer. After purification, 5’ Illumina TruSeq sequencing adapters were ligated to the cDNA. After PCR amplification, the amplified cDNA was purified using the Agencourt AMPure XP kit (Beckman Coulter Genomics) and evaluated via bioanalyzer. The cDNA was pooled and further purified (200-600 bp) using a preparative agarose gel. The finished pooled libraries were sequenced by the Core unit SysMed (University of Würzburg) using an Illumina NextSeq 500 system and 75 bp read length.

Reads from the RNA-seq were trimmed and filtered using the FASTX toolkit (v.0.10.1; http://hannonlab.cshl.edu/fastx_toolkit). Mapping was performed using READemption (v.1.01) against the genome sequence for *F. nucleatum* subsp. *nucleatum* ATCC 23726 (NZ_CP028109.1) with custom annotation. Differential gene expression analysis was performed using DEseq2 (v.1.18.01). The data from three replicates was used for the analysis and the genes showing a log_2_ fold change ≤ -2 or ≥ 2 with false-discovery-rate (FDR) was ≤ 0.05 are considered as differentially regulated genes.

### Rifampicin assay

*F. nucleatum* WT and deletion strains (Δ*khpA*, Δ*khpB*, Δ*khpA*Δ*khpB*) were grown to mid-exponential phase (OD_600_=0.5) and rifampicin was added to a finial concentration of 240 μg/ml. A volume of 10 ml cells was collected immediately, or different time points after adding rifampicin (3, 6, 12, 24, 48 min) in a 15 ml falcon tubes containing 2 ml stop mix. The samples were snap frozen in liquid nitrogen and stored at -80 °C, further subjected to RNA extraction by hot phenol and DNase I digestion. 5 μg of each DNase I-digested RNA sample was loaded on 6% acrylamide gels containing 8 M urea and subjected to northern blot analysis as described above. The RNA stability was determined by measuring band intensities with ImageJ and calculating half-lives using Prism.

### RNA-coimmunoprecipitation

RNA coimmunoprecipitation was performed as previously carried out for Hfq or ProQ (33,37). Briefly, bacterial cultures (4 replicates) were grown to mid-exponential phase (OD_600_= 0.5) and a volume of cells equal to 100 ODs was collected. Cells were suspended in 800 μl lysis buffer (20 mM Tris pH 8.0, 150 mM KCl, 1 mM MgCl_2_,1mM DTT) containing 8 μl of DNase I. The resuspended cells were added to a fresh 2ml tube with 800 μl of 0.1 mm glass beads and lysed at 30 Hz for 10 min using Retsch MM200. A volume of supernatant equivalent to 0.5 OD_600_ cells was added to 90 μl 1x protein loading dye and used for input control. 25 μl of Flag antibody (Monoclonal ANTI-FLAG M2, Sigma, #F1804) was added to the supernatant and rotated for 30 min at 4 °C. The sample was then added with 75 μl pre-washed Protein A Sepharose (Sigma, #P6649) and incubated at 4 °C for 30 min with rotation. Afterwards, the beads were spun down and a volume equivalent to 0.5 OD_600_ cells were added to 90 μl 1x protein loading dye and used for flowthrough control. The beads were then washed 5 times with the lysis buffer by inverting the tubes and centrifuged roughly. The last washing was collected in 1x protein loading dye. After washing, resuspended the beads in 532 μl lysis buffer and collected 32 μl to 8 μl 5x protein loading dye for later western blot analysis. The remaining beads were added 500 μl P: C: I for RNA precipitation. After precipitation, the RNA pellets were resuspended in H_2_O and further subjected to DNase I digestion and purified by P: C: I. At the end, the RNA pellets were resuspended in 15 μl H_2_O and used 15 μl from one replicate for northern blot verification and 3.5 μl of other three replicates for the cDNA library preparation.

### cDNA library preparation

For preparation cDNA library of the immunoprecipitated RNA samples, NEBNext Multiplex Small RNA Library Prep Set for Illumina (New England Biolabs) was used. Briefly, 3.5 μl of RNA was mixed with 1 μl of 3’ SR Adaptor (diluted 1:10 in nuclease-free water) and incubated in a preheated thermal cycler for 2 min at 70 °C. The samples were transferred to ice and added with the premixed 3’ ligation mix (5 μl 3’ Ligation Reaction Buffer, 1.5 μl 3’ Ligation Enzyme Mix). The ligation mixture was then incubated for 1 h at 25 °C. 0.25 μl of SR RT Primer and 2.5 μl nuclease-free water were added to the mixture and incubated at 75 °C for 5 min, 37 °C for 15 min and followed by 25 °C for 15 min. To ligate the 5’ SR adaptor, the 5’ ligation mixture (0.5 μl 1:10 diluted 5’ SR Adaptor, 0.5 μl 10x 5’ ligation reaction buffer, 1.25 μl 5’ ligation enzyme mix) was added to each sample and incubated for 1 h at 25 °C. For synthesis first strand of cDNA, reverse transcription mix (4 μl first strand synthesis reaction buffer, 0.5 μl murine RNase inhibitor, 0.5 μl M-MLV reverse transcriptase) was added to each reaction and incubated for 1 h at 50 °C followed by 15 min at 70 °C to inactivate the reverse transcription reaction. 5 μl of cDNA was then proceeded to PCR amplification with the PCR mix (12.5 μl LongAmp Taq 2x master mix, 0.625 μl SR primer, 6.25 μl nuclease-free water) and 0.625 μl index primer. PCR program (30s at 94°C, 15 s at 94 °C, 30 s at 62 °C and 15 s at 70 °C for 19 cycles, 5 s at 70 °C) was carried out. PCR products were mixed with 25 μl of GLII buffer and loaded on 6 % polyAcrylamide gel containing 8 M urea. After staining the gel for 10 min in 1x TBE buffer with a 1:10,000 dilution of SYBR Gold Nucleic Acid Gel Stain (Invitrogen), the bands in the range of 130- 1000 bp were cut into small pieces and transfer to a 2 ml LoBind tube. 500 μl of DNA elution buffer was added and incubated at the room temperature for 2 h. After transferring the elute and the gel debris to centrifuge tube filters (Sigma) and centrifugation (5000 rpm, 1 min), the elute was recovered by adding 1 μl liner acrylamide (NEB) and precipitating in ethanol at -80°C for 1 h. After precipitation, the pellet was washed with 80% ethanol and resuspended in 12 μl nuclease free water (NEB). 1 μl of the prepared cDNA library was subjected to Bioanalyzer analysis.

Amplified cDNAs were pooled and sequenced on an illumine NextSeq 2000 platform by the CoreUnit Sysmed from the University of Würzburg.

### RIP-seq data analysis

Approximately 40 million paired-end 75 bp reads were sequenced for each individual cDNA library. Generated FASTQ files were mapped to *F. nucleatum subsp. nucleatum* ATCC 23726 (NC_003454.1) genome with custom annotation. Gene-wise read counts were normalized to TPM (transcripts per kilobase million) that takes into account sequencing depth (number of mapped reads) and transcripts length. Enrichment factors were calculated with DESeq2. RNA coimmunoprecipitated with tagged KhpA-3×FLAG or KhpB-3×FLAG (three biological duplicate each) was compared to the non-tagged WT strain to determine enrichment factors. DESeq2 utilizes Wald test to determine the *P*-value and the Benjamini-Hochberg to adjust P-values (P- adj). Transcripts enriched with log_2_ fold change ≥ 2.0 with *P*-adj ≤ 0.05 were considered binding ligands of KhpA or KhpB (Supplementary Tables S2).

### Western blot

For the protein samples from the RNA affinity purification and RNA immunoprecipitation, proteins were loaded on 15% SDS-PAGE gel and transferred to PVDF filter and immunodetection.

As primary antibody, the anti-KhpB was generated with a synthesized peptide derived from C-terminus of KhpB protein (H-CGRDPKRYIVIKKKRG-OH) in rabbits (Eurogentec). The KhpB anti- serum was tested with ELISA assay and further cleaned with affinity purification, stored in 1x PBS with 0.01% thimerosal and 0.1% BSA. 1: 200 dilution was used for KhpB detection. The monoclonal anti-FLAG M2 (Sigma, #F1804) was 1: 1000 diluted for detecting FLAG-tagged KhpA or KhpB. The anti-mouse (Thermo Fisher# 31430, 1: 5000 diluted) and anti-rabbit (Thermo Fisher# 31460, 1: 10000 diluted) conjugated to horseradish peroxidase were used as secondary antibodies for anti-KhpB and anti-FLAG respectively. For detection, ECL™ Prime Western Blotting Detection Reagent (GE Healthcare) used as a substrate.

### Microscopy assay and cell length measurements

*F. nucleatum* cells were cultured to mid-exponential phase (OD_600_=0.5) with three biological replicates. Cultures were collected and pellets were washed with 1xPBS and resuspended in 1x PBS, further subjected to fix with 4% PFA for 20min at 4 °C. After fixing, the cells were further stained with DAPI and FM 4-64. The samples were imaged on agarose chambered coverslips with a Leica SP5 laser scanning confocal microscope (Leica Microsystems).

For measuring the cell length, the cells with a single DAPI-staining spot were evaluated with ImageJ manually. Visibly separated cells or the cells with two or more DAPI-staining spots were excluded.

### eut genes expression analysis by qRT-PCR

For *eutS* and *eutL* gene expression level analysis, *F. nucleatum* WT, deletion (Δ*khpA*, Δ*khpB*, Δ*khpA*Δ*khpB*) and complementary strains were grown to late-exponential phase (OD_600_= 0.8) in Columbia broth. The cells were spun down (4,000 x*g* for 3 min) and washed with 1xPBS, transferred into CYG medium complemented with 25 µM ethanolamine or CYG medium, separately. After re-culturing the cells for 2 h, RNA samples were collected. Total RNAs were extracted with Trizol (Invitrogen) and followed by DNase I (ThermoFisher Scientific) digestion. cDNAs for qRCP were generated with 1 µg DNase treated total RNA samples using the M-MLV reverse transcriptase and random hexamer primers (ThermoFisher Scientific) following the manufacturer’s instructions. The synthesized cDNAs were diluted at 1: 100 and subjected to qPCR using Takyon Mater Mix (Eurogentec) on CFX system (Bio-Rad). 5S RNA served as reference control. Oligonucleotides used are listed in Supplementary Table S4. The qPCR data was analyzed using the double delta Ct analysis (38).

## RESULTS

### An RNA-centric approach to identify RNA-protein interactions in F. nucleatum

To identify RNA-protein interactions in *F. nucleatum*, we initially explored the widely used MS2 aptamer approach (39) to tag sRNAs from *F. nucleatum*, but with limited success. Therefore, we used the 14-mer capture tag to isolate sRNA-associated RBPs in this species. We selected 6S RNA to establish proof-of-concept of this method in *F. nucleatum*. 6S RNA is a sRNA regulator of transcription broadly present in bacteria and it forms a stable complex with RNA polymerase (RNAP) (40,41). RNAP can liberate itself from 6S RNA by using the latter as a template for RNA synthesis, producing tiny RNA products (pRNAs) (42). We have previously detected pRNAs in *F. nucleatum* (7), predicting the formation of a 6S RNA-RNAP complex in this bacterium.

We attached the 14-mer capture tag to the 5’ end of 6S RNA, *in vitro* transcribed the construct and recovered the adaptor coupled 6S RNA after incubation with an *F. nucleatum* lysate using Streptavidin beads (Figure 1B). The experiment was performed using a bacterial strain that expresses a FLAG-tagged version of the alpha subunit of RNAP (RpoA-3xFLAG) from a plasmid and we were able to confirm recovery of RpoA by western blot analysis (Figure 1B). The sequence tag alone, which served as a negative control, did not recover RpoA (Figure 1B).

Next, we analyzed the elution samples of the pulldown experiment using a lysate from a *F. nucleatum* WT strain by mass spectrometry and observed enrichment for all core RNAP components (RpoA, RpoB, RpoC, RpoZ) and for the sigma factor RpoD (Figure 1C, Supplementary Table S1). In addition to the bona fide RNAP components, we detected enrichment of the small ribosomal subunit protein RpsO (C4N14_03125), DNA gyrase subunit A, GyrA (C4N14_02325), and a protein of unknown function (C4N14_02505), indicating that these proteins interact with the 6S RNA or 6S RNA-RNAP complex in *F. nucleatum*.

Together, these data establish an RNA-centric strategy to uncover RNA-protein interactions in *F. nucleatum* and suggest that 6S RNA-RNAP complex formation is conserved in this bacterium.

### The landscape of sRNA-associated RBPs in F. nucleatum

To systematically identify sRNA-associated RBPs in *F. nucleatum*, we *in vitro*-synthesized 19 additional tagged sRNAs that covered four classes of recently identified fusobacterial sRNAs (intergenic, antisense, 5’ UTR-derived and 3 ’UTR-derived) (7). The tagged sRNAs were used as baits in pulldown experiments with *F. nucleatum* lysates from mid-exponential phase when the bacterium expresses most of its sRNAs (7). The elution fractions were subjected to MS. Across all sRNAs, we identified a total of 75 proteins that were significantly enriched compared to the control (log_2_ ratio ≥ 4 and log_10_ LFQ intensity ≥ 5) (Supplementary Fig. 1, Supplementary Table S1). 24 of these 75 proteins were classified as ‘RNA-binding’ according to Gene Ontology (GO) terminology (43) and included 16 ribosomal proteins, 4 putative ribonucleases (RNase III, RNase J, RNase R, PNPase), the ATP-dependent RNA helicase DEAD, and 3 predicted RBPs (C4N14_06085, C4N14_04780, C4N14_02375). In addition, we identified translation initiation factors (2/75), RNA polymerase subunits (2/75), a DNA-binding protein (1/75), and a cold shock family protein (1/75). Proteins of unknown function (14/75) were also prominent in the sRNA pulldowns (Supplementary Table S1) and included proteins with interesting protein domains, such as the S1 RNA binding domain protein C4N14_06955.

To further classify these proteins, we examined their COG (clusters of orthologous groups) categorization (44). Consistent with the frequent enrichment of ribosomal proteins, we found that 24 of the 75 enriched proteins belonged to the COG group linked to ‘translation, ribosomal structure and biogenesis’ (Supplementary Fig. 1). We also observed enrichment for COGs associated with metabolism (Supplementary Fig. 1). Proteins that fall into these clusters include the acyl carrier protein AcpP (C4N14_04130), which is involved in fatty acid biosynthesis and a deblocking aminopeptidase involved in carbohydrate transport and metabolism (C4N14_07180). This observation is in line with studies that identified metabolic proteins as putative RBPs (45,46).

Since we were interested in global sRNA binding proteins, we searched for proteins that interacted with multiple sRNAs. The top candidates included C4N14_04780 and C4N14_02375, two putative homologs of the RBPs KhpA and KhpB (16/19 baits), a histone-like HU family DNA-binding protein (C4N14_08125) (15/19 baits), AcpP (15/19 baits), the ribosomal protein RpsK (C4N14_05705) (14/19 baits) and a putative cold shock protein C4N14_09590 (14/19 baits) (Figure 2 and Supplementary Table S1).

**Figure 2.**
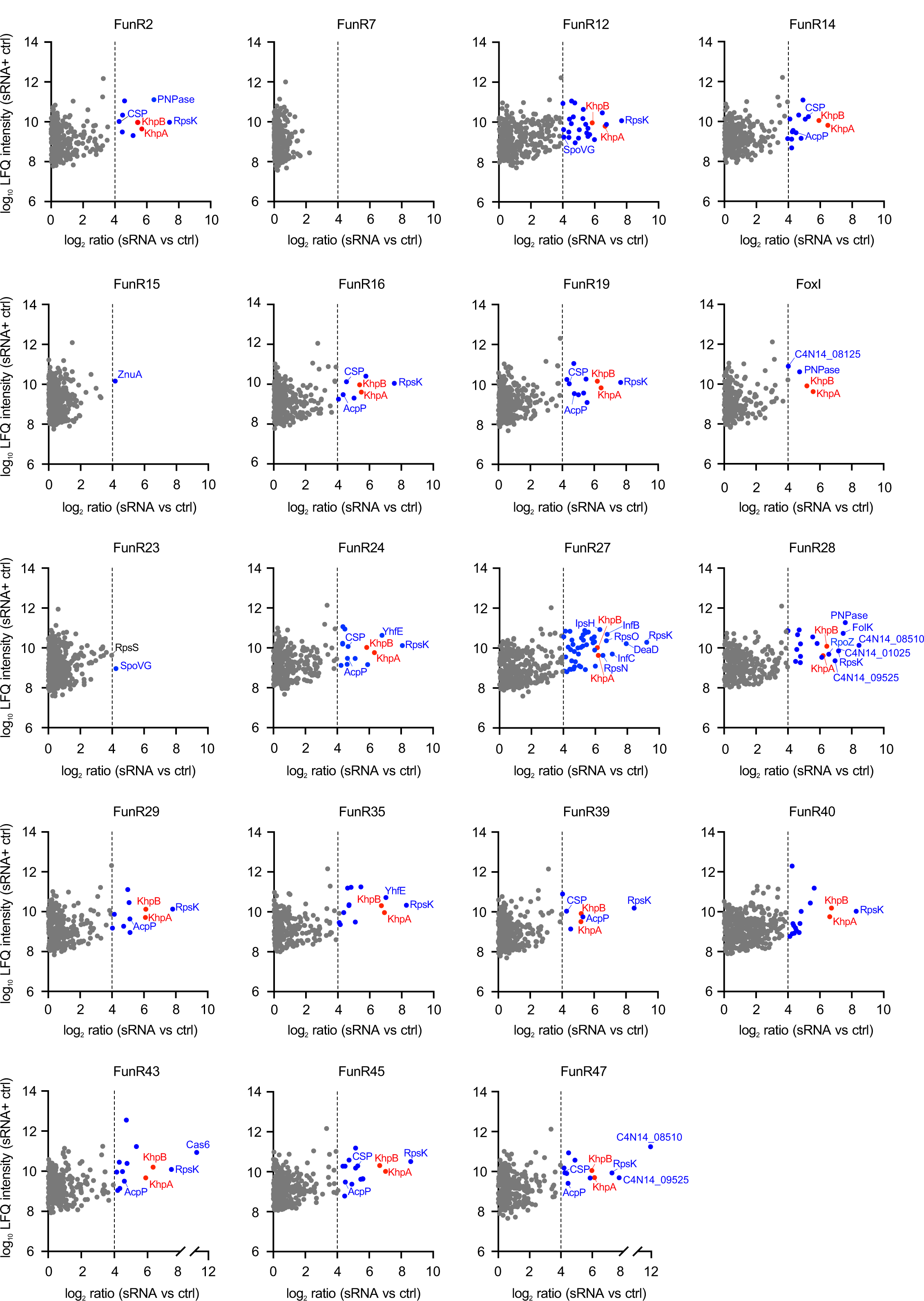
The landscape of sRNA-associated RBPs in *F. nucleatum*. Mass spectrometric analysis of sRNA affinity purifications with 19 sRNA baits. Log_10_ LFQ intensities of the bait sRNAs and the control RNA are plotted against the log_2_ ratios of the sRNAs versus the control. Cut-off for significantly enriched proteins (blue or red dots) was set to 4 log_2_ ratios (dotted line). Interesting protein candidates are labeled.

Thus, our affinity purification approach led to the identification of both specific and broad interactors of the different *F. nucleatum* sRNAs.

### KhpA and KhpB in F. nucleatum show conserved domain organization and genomic synteny

Given the recovery of KhpA and KhpB with many different sRNAs in *F. nucleatum* (Figure 3A) and their recent identification as global RBPs in several Gram-positive bacteria such as *S. pneumoniae*, *Clostridioides difficile*, and *Enterococcus faecalis* (34–36), we decided to focus on these two RBP candidates for further analysis. Both KhpA and KhpB belong to the K-homology (KH) domain-containing protein family, which is prevalent in different phyla, such as Actinobacteria and Firmicutes (47). The KH domain is a well-characterized RNA-binding domain that contains a conserved GXXG motif, which interacts with RNA backbone residues (48). KhpA is a single KH domain containing protein, while KhpB has an N-terminal Jag-N domain of unknown function, followed by a linker region that connects to the KH domain and a second R3H RNA-binding domain, which harbors a RXXXH RNA-binding motif (34). In *F. nucleatum*, this domain composition of both proteins is conserved (Figure 3B).

**Figure 3.**
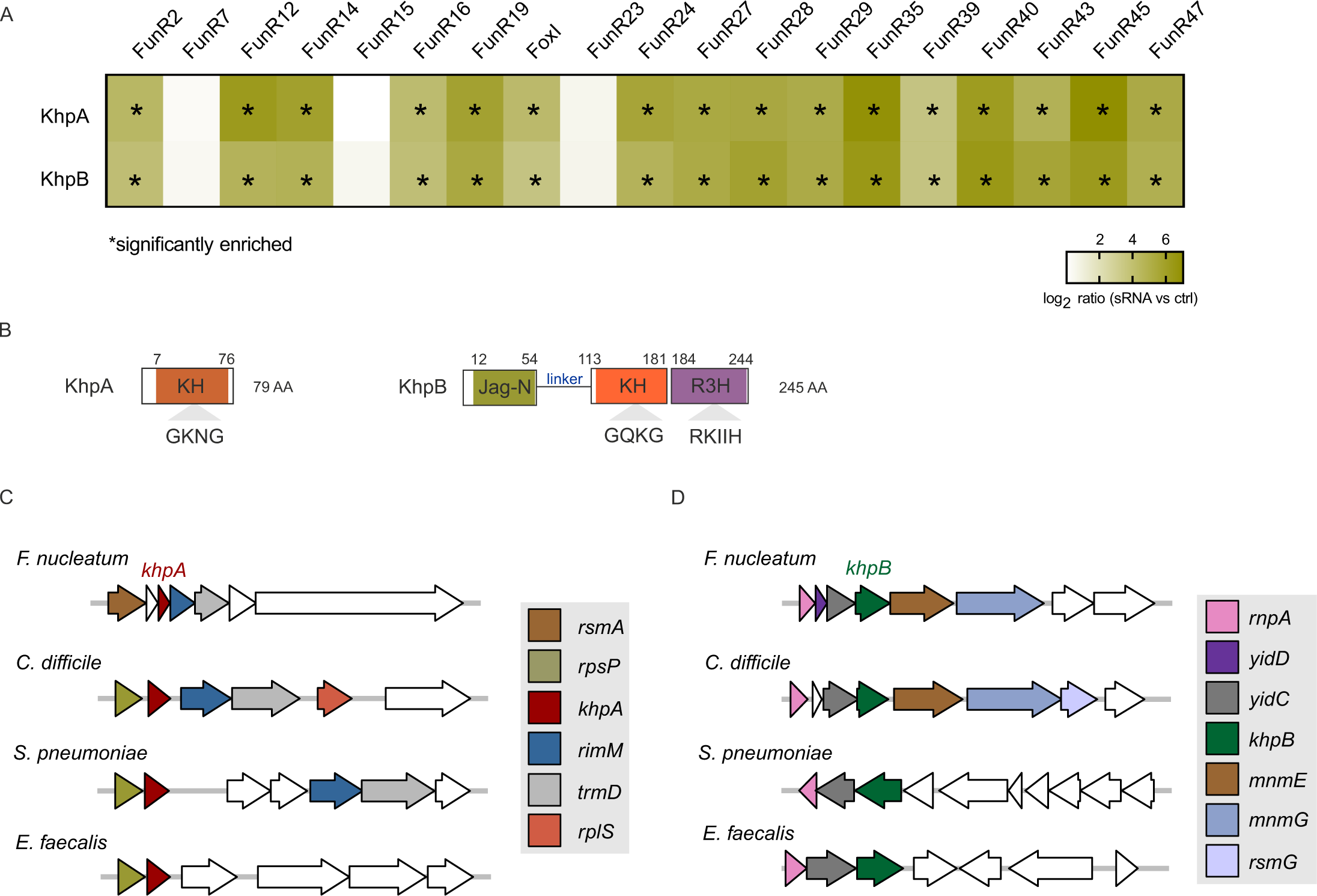
KhpA and KhpB proteins. (**A**) Heat map of the log_2_ ratios of sRNAs versus control pull-downs for KhpA and KhpB based on the mass spectrometric data. Asterisks indicate significant enrichment (log_2_ ratio ˃4). (**B**) General domain composition of KhpA or KhpB. The KH (*K- homology*) domains are colored orange in KhpA and KhpB proteins,whereas the KhpB Jag-N and R3H domains are colored in green and purple, respectively. (**C-D**) Gene synteny surrounding *khpA* (C) or *khpB* (D) in *F. nucleatum* and three Gram-positive bacteria (*S. pneumoniae*, *C. difficile*, *E. faecalis*). The gene sequences were obtained from KEGG and gene synteny was visualized with the R package gggenes.

Despite the conserved domain organization, the amino acid sequence conservation of *F. nucleatum* KhpA and KhpB and their respective homologues in other bacterial species is low (≤ 33 % identity for KhpA and ≤ 35% for KhpB) (Supplementary Fig. 2A and S2B). Especially the linker region between the Jag-N and KH domain in KhpB varies greatly (Supplementary Fig. 2B). Nevertheless, the RNA binding motifs (CXXG in KhpA and KhpB, RXXXH in KhpB) are conserved (Supplementary Fig. 2A and S2B). In contrast to the lack of sequence conservation among KhpA and KhpB homologs in general, KhpA and KhpB show strong amino acid sequence conservation across all *F. nucleatum* species (≥ 91% for KhpA and ≥ 80% for KhpB) as well as in the related species *F. periodonticum* and *F. hwasookii* (Supplementary Fig. 3A and S3B).

As noted previously in different Gram-positive bacteria (35,47), we observed that the genes flanking *khpA* and *khpB* show conserved genomic synteny in *F. nucleatum* (Figure 3C and 3D). The *khpA* gene is co-conserved with *trmD*, which encodes a tRNA methyltransferase and the 16S rRNA processing protein *rimM* (Figure 3C). The *khpB* gene is part of an operon that includes *rnpA*, which encodes the RNA component of ribonuclease P and the membrane protein insertase *yidC* (Figure 3D).

Overall, their domain composition, sequence conservation of the RNA-binding motifs, and gene synteny suggests a functional conservation of KhpA and KhpB in fusobacteria.

### KhpB binds directly to sRNAs

To validate our global MS data, we repeated the sRNA pulldown for the two sRNAs FoxI and FunR2, both of which showed strong interaction with KhpA and KhpB in our initial data set (Figure 2). Western blot analysis of FoxI and FunR2 pulldowns using a KhpB antibody confirmed the recovery of KhpB; whereas no signal was obtained with the sequence tag alone, included as a control (Figure 4A).

**Figure 4.**
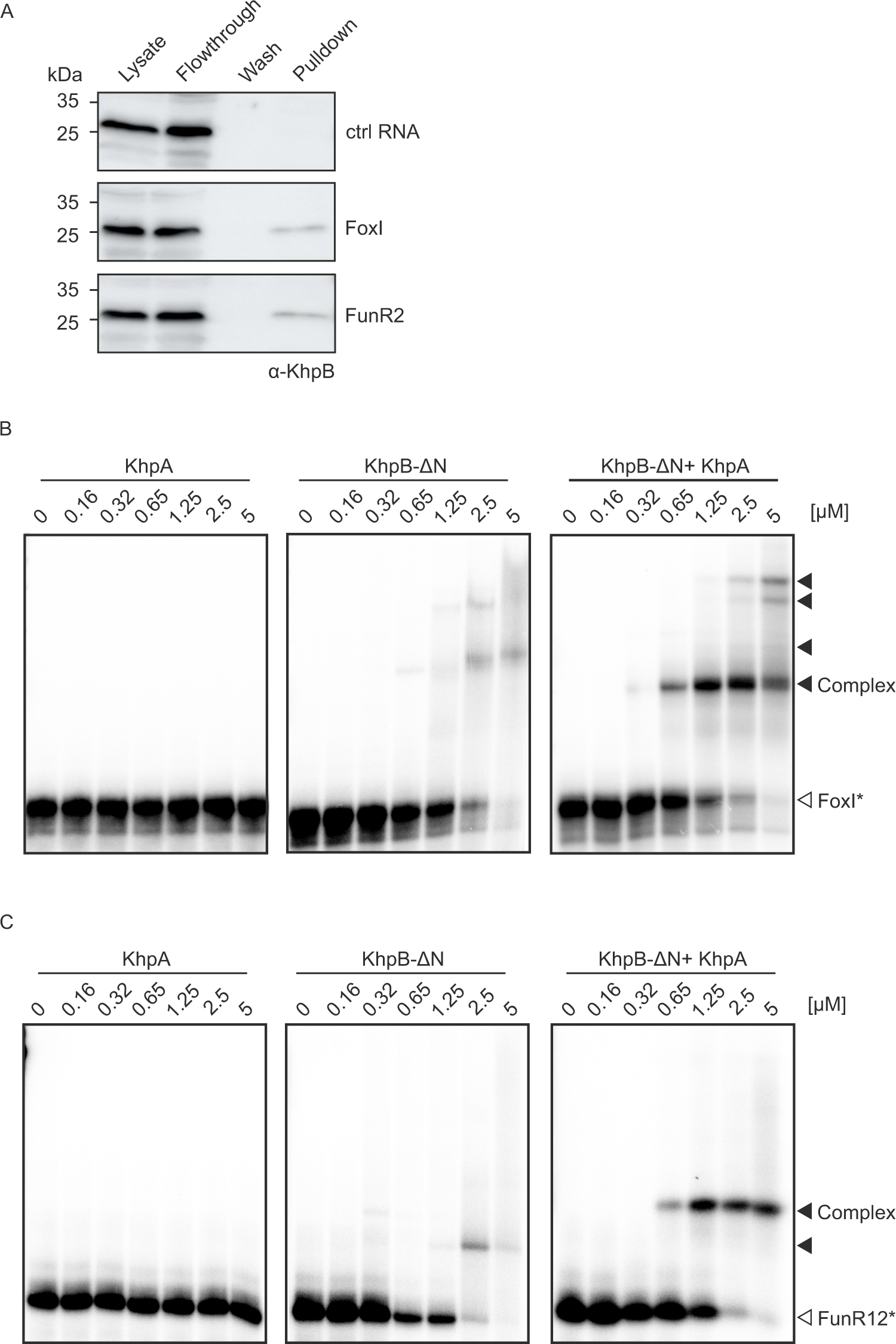
KhpB binds directly to sRNAs. (**A**) Western blot analysis of protein samples collected from different steps of the sRNA affinity purification performed with the control RNA, tagged FoxI or tagged FunR2 using anti-KhpB antiserum. (**B-C**) EMSAs using purified KhpA, KhpB-ΔN, or both proteins (1:1 ratio) with FoxI (B) and FunR12 (C). 4 nM of *in vitro* transcribed and radiolabeled FoxI or FunR12 were incubated with increasing concentration of KhpA, KhpB or both proteins (0, 0.16, 0.32, 0.63, 1.25, 2.5, 5 µM).

To reconstitute complex formation *in vitro*, we sought to purify recombinant *F. nucleatum* KhpA and KhpB proteins expressed in *E. coli*. We were able to isolate soluble KhpA, but full-length KhpB was insoluble. Therefore, we used a truncated version of KhpB (KhpB-ΔN, aa 83-245) that lacks the Jag-N domain but still contains the KH and R3H domains, which are important for RNA-binding. We purified full-length KhpA and KhpB-ΔN via a cleavable 6xHis-tag, followed by ion-exchange chromatography after 3C protease cleavage to remove the tag (Supplementary Fig. 4).

To test if the interaction between KhpA/KhpB and FoxI or FunR2 is direct, we performed electrophoretic mobility shift assays (EMSAs). Incubation of both sRNAs with KhpA did not lead to a shift, even at the highest concentrations tested (Figure 4B and 4C, left panel). Incubating radioactively-labeled FoxI and FunR12 with increasing concentrations of KhpB-ΔN, we observed a weak shift, indicative of an interaction between KhpB and the two sRNAs. Since prior studies in *S. pneumoniae* suggested that KhpA and KhpB act as heterodimer to bind RNA (34), we tested if co-incubation with both proteins would increase the RNA-binding affinity of KhpB and lead to the formation of a distinct complex. In fact, we observed much stronger binding of both sRNAs when performing the EMSAs with KhpA and KhpB together (Figure 4B and 4C, right panel). We also detected the formation of different RNA-protein complexes, suggesting that either KhpA or KhpB form multimers or that they bind multiple copies of the sRNAs. Overall, these results indicate that KhpB directly binds to sRNAs and that this activity is enhanced by KhpA, supporting the role of KhpA and KhpB as sRNA-associated RBPs in *F. nucleatum*.

### KhpA and KhpB influence the stability of sRNAs

One frequently observed consequence of a specific association between a bacterial sRNA with a global RBP is an effect on transcript stability; for example, many sRNA targets of ProQ show altered (primarily reduced) intracellular stability in a *proQ* deletion strain of *Salmonella* (33). Therefore, we wondered if KhpA and KhpB affect the stability of sRNAs in *F. nucleatum*. To address this question, we first generated single in-frame deletion mutants of *khpA* (Δ*khpA*), *khpB* (Δ*khpB*) and a double deletion strain (Δ*khpA*Δ*khpB*) using a recently developed gene deletion system (Supplementary Fig. 5A) (5).

We then used these strains to investigate the effect of KhpA and KhpB on RNA stability using rifampicin to arrest bacterial transcription. After adding rifampicin to the bacterial cell culture, RNA samples were collected at different time points and subjected to northern blot analysis to determine RNA half-lives. We observed that the half-life of FoxI is reduced from ∼3.8 min in the wild type (WT) strain to <2 min in all deletion strains (Δ*khpA*, Δ*kphB*, Δ*khpA*Δ*kphB*) (Figure 5A and 5B). The lack of *khpA* or *khpB* also reduced the stability of FunR12 (Figure 5A and 5C). In both cases, the deletion of *khpA* had a weaker effect compared to deletion of *kphB* and the double-deletion (Δ*khpA*Δ*kphB*) (Figure 5A-C). Interestingly, we also observed that the deletion of *khpA* or *khpB* resulted in an increased stability of the sRNA FunR2. The half-life of FunR2 doubles to 8 min in Δ*khpA* or Δ*kphB* and ∼9.5 min in Δ*khpA*Δ*kphB*, as compared to ∼4 min in the WT strain (Figure 5A and 5D). These data indicate that KhpA and KhpB can affect sRNA stability both positively and negatively.

**Figure 5.**
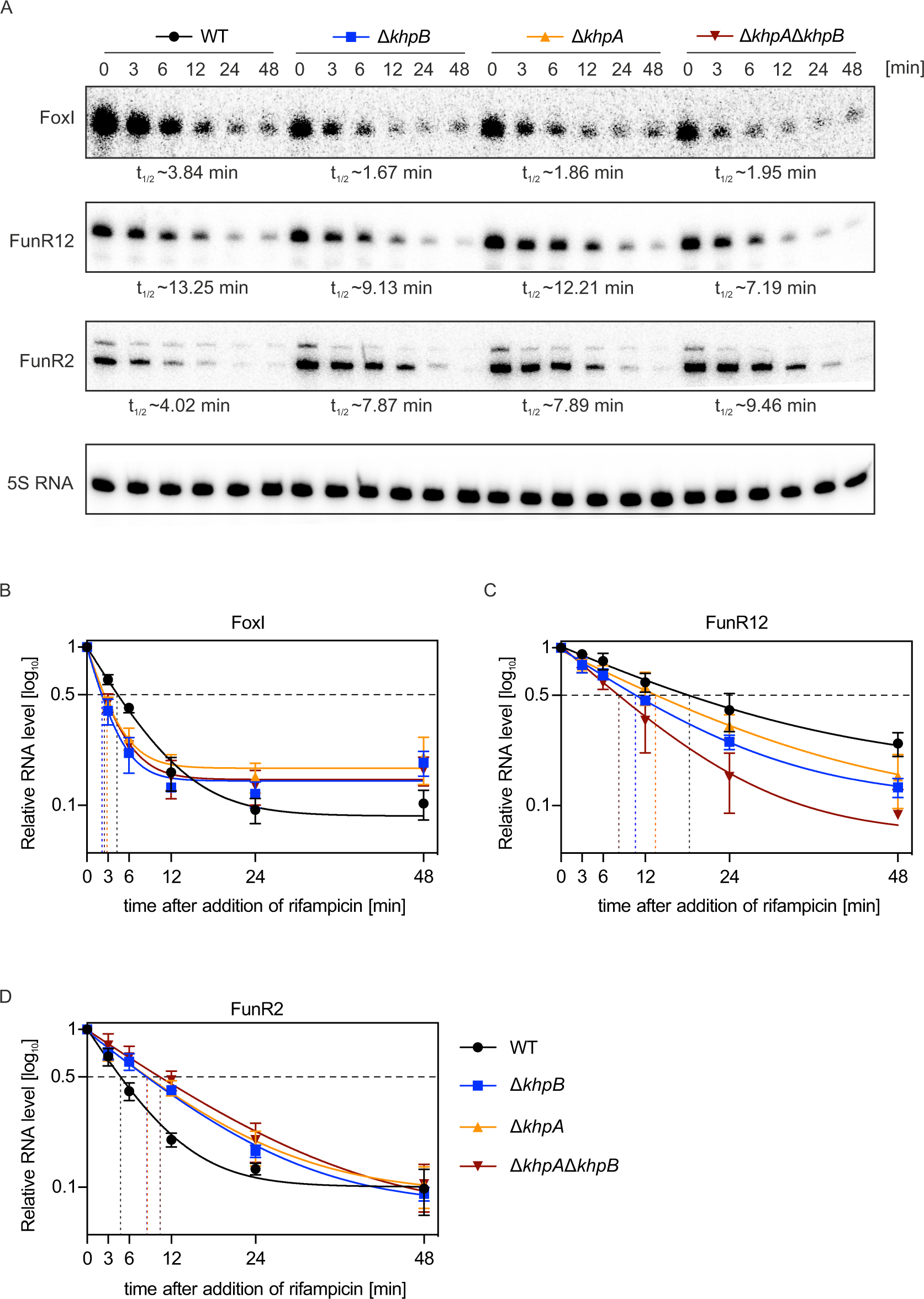
KhpA and KhpB influence the stability of sRNAs. (**A**) Stability of FoxI, FunR12 and FunR2 in WT, Δ*khpA*, Δ*khpB*, Δ*khpA*Δ*khpB* strains determined by detection of RNA abundance via Northern blot after rifampicin treatment. Samples were collected at 0, 3, 6, 12, 24, 48 min after addition of rifampicin. 5S RNA was probed as loading control. The half-life of each sRNA in each strain is indicated beneath each blot. The blot is a representative of three experimental replicates. (**B-D**) Quantifications of FoxI (B), FunR12 (C) and FunR2 (D) decays. For quantification, band intensities of the blots (A) were measured with ImageJ and half-lives were calculated with GraphPad Prism. Mean value ± SD from three replicates is shown.

### KhpA and KhpB are global RBPs

To define the full target spectrum of KhpA and KhpB, we performed RNA immunoprecipitation followed by deep sequencing (RIP-seq) to capture transcripts that interact with KhpA and KhpB. As a first step, we generated chromosomally 3xFLAG-tagged versions of KhpB and KhpA. The *khpA^3xFLAG^* was generated in the native locus (Supplementary Fig. 5B). Since we observed a growth defect in the *khpB^3xFLAG^* strain, we placed KhpB-3xFLAG under a constitutive promoter in an insulated complementation locus in the Δ*khpB* background (Supplementary Fig. 5B). Western blot analysis confirmed similar expression levels of KhpB in the WT and Δ*khpB^khpB-3xFLAG^* strain in the mid-exponential phase (Supplementary Fig. 5C). Both tagged strains show similar growth characteristics to the WT strain (Supplementary Fig. 5D).

We then performed RIP-seq in the *khpA^3xFLAG^*, Δ*khpB^khpB^*^-*3xFLAG*^ and WT strain. Western blot analysis confirmed recovery for both KhpA and KhpB, although KhpA was recovered less efficiently (Figure 6A, upper panel). RNA-seq of the immunoprecipitated RNAs shows distribution of mapped reads across all RNA classes (5’-UTRs, CDSs, 3’-UTRs, rRNAs, tRNAs, sRNAs) in the WT, *khpA^3xFLAG^* and Δ*khpB^khpB^*^-*3xFLAG*^ strain. Reads mapped to CDSs showed a higher percent of total reads in KhpA and KhpB pulldowns compared to WT control (Figure 6B, Supplementary Table S2). DEseq2 analysis revealed 151 transcripts that are significantly enriched by KhpA and 562 transcripts by KhpB (log_2_ fold change ≥ 2.0, *p-adj* ≤0.05), and the reads mapped to 5’-UTRs, CDSs, 3’-UTRs, tRNAs, as well as sRNAs (Figure 6C, Supplementary Table S2). A comparison of the different RIP-seq data sets showed that the majority of transcripts bound by KhpA (105/151) were also enriched by KhpB (Supplementary Fig. 5E, Supplementary Table S2). This further supports the notion that both proteins cooperate in RNA-binding, which is consistent with their homologues in *S. pneumoniae* (34,49). Among the significantly enriched transcripts that were found in both data sets were the 3’-UTRs of the transcripts encoding the cell division proteins FtsL and FtsZ (Figure 6C). Interestingly, mRNAs encoding cell division proteins were also enriched in KhpB RIP-seq data in *S. pneumoniae* and *C. difficile* (34,35). We also noticed that certain tRNAs enriched by KhpA (tRNA^Val^) and KhpB (tRNA^Cys^, tRNA^Asn^, tRNA^Tyr^) in *F. nucleatum* also interact with KhpB in *S. pneumoniae* and *E. faecalis* (34,36) (Supplementary Table S2). While the biological function of these interactions is unknown, our data indicate that KhpB has multiple conserved binding partners.

**Figure 6.**
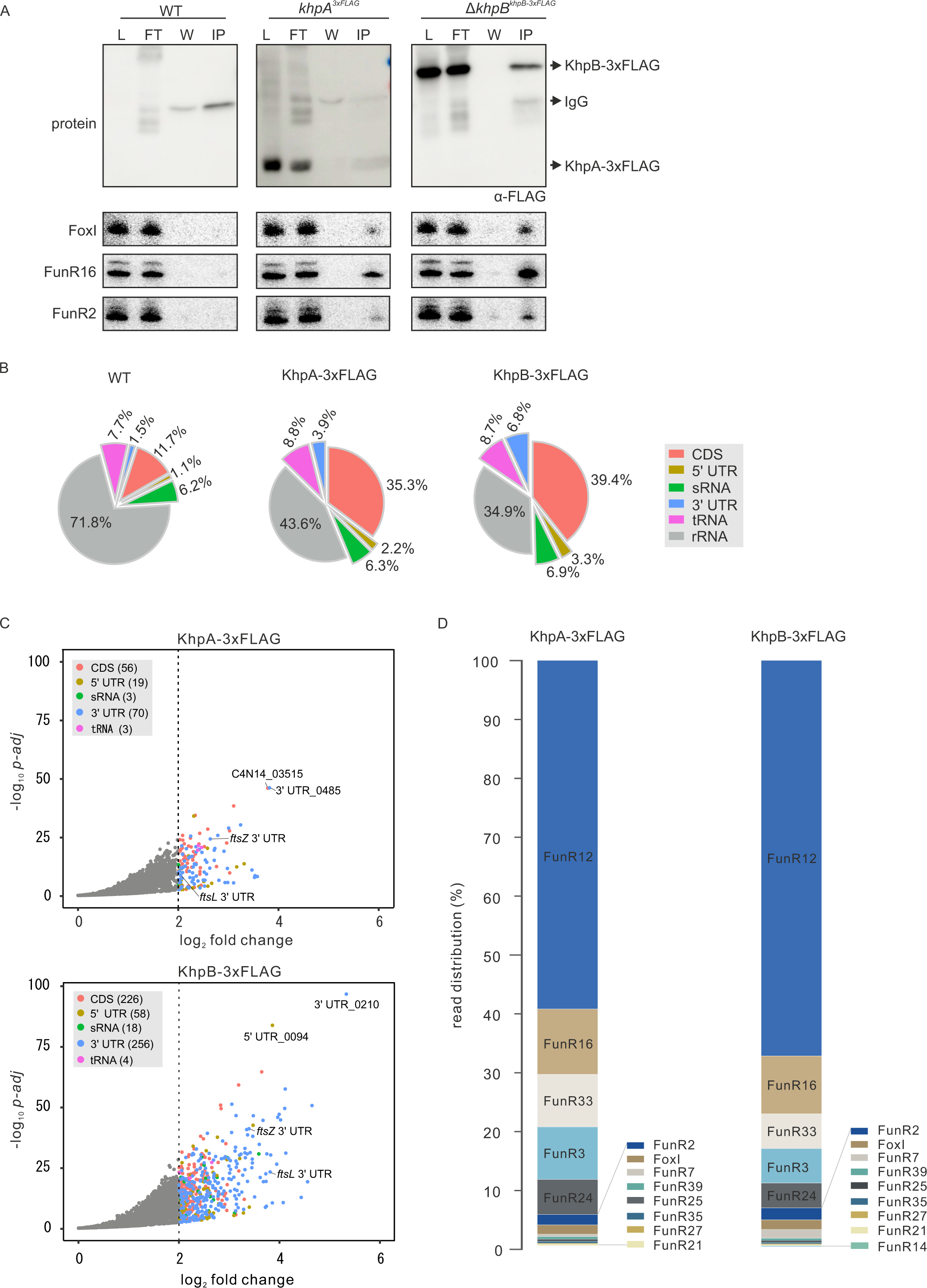
KhpA and KhpB are global RBPs. (**A**) Detection of immunoprecipitated proteins (upper panel) or RNAs (lower three panel) from RIP experiment. Upper panel: KhpA-3xFLAG or KhpB-3xFLAG was detected on western blot with a FLAG-specific antibody in lysate (L), flow-through (FT), wash (W) and immunoprecipitation fractions (IP). Lower panel: Northern blot detection of FoxI, FunR16 or FunR2. The WT strain was used as a negative control. (**B**) Pie charts representing the read distribution of the indicated RNA classes in WT, KhpA-3xFLAG and KhpB-3xFLAG RIP-seq. (**C**) Volcano plots of RNA transcripts enriched by KhpA-3xFLAG and KhpB-3xFLAG in RIP-seq. DEseq2 was used to determine the enrichment factors between FLAG-tagged strains and the WT control. Cut-off (dashed line) for significantly enriched transcripts was set to a 2 log_2_ fold-change. Significantly enriched transcripts from different RNA classes are highlighted in color. Numbers in parentheses give the number of significantly enriched transcripts belonging to the respective RNA class. (**D**) Chart showing read distribution of sRNAs associated with KhpA-3xFLAG and KhpB-3xFLAG in RIP-seq. Percentage indicates the reads of a sRNA compared to all sRNAs in a cDNA library.

In line with our initial sRNA affinity purification experiments, we observed that sRNAs were recovered by KhpA and KhpB. The highest read distribution is seen for the sRNA FunR12, with 59% of all sRNA mapped reads for KhpA and 62% for KhpB (Figure 6D). Other highly enriched sRNAs include FunR16, which contributes 10% of all sRNA reads for KhpA or KhpB, respectively (Figure 6D). Northern blot analysis confirmed the enrichments of several selected sRNAs (FoxI, FunR2 and FunR16) in the RIP experiment. Again, KhpB showed higher apparent binding affinity for these three sRNAs compared to KhpA (Figure 6A, lower panel). Taken together, our RIP-seq data sets indicate that KhpA and KhpB not only bind multiple sRNA but also act as global RBPs in *F. nucleatum*.

### KhpA and KhpB regulate gene expression

The interaction of KhpA and KhpB with a wide variety of RNAs in *F. nucleatum* suggests a global effect on gene expression. We therefore compared the steady-state RNA levels between WT *F. nucleatum* and the individual deletion strains (Δ*khpA* or Δ*khpB*) by RNA-seq analysis in the mid-exponential and early-stationary growth phase. In the mid-exponential phase, 27 transcripts were differentially expressed in Δ*khpA* and 116 in Δ*khpB* (log_2_ fold change ≥ 1.0, *p-adj* ≤0.05 or log_2_ fold change ≤ -1.0, *p-adj* ≤ 0.05) (Figure 7A, Supplementary Table S3). In the early-stationary phase, both Δ*khpA* and Δ*khpB* regulated a larger suite of genes, with 117 and 179 differentially expressed transcripts in Δ*khpA* and Δ*khpB*, respectively (Figure 7B, Supplementary Table S3). An operon of unknown function (C4N14_09375-C4N14_09395) shows the highest upregulation in the absence of KhpA and KhpB in both growth phases (Figure 7A-B). In the stationary phase, general stress operons (*hrcA*-*grpE*-*dnak*-*dnaJ* and *groELS*) as well as molecular chaperones (*htpG* and *clpB*) are highly up-regulated in the Δ*khpA* and Δ*khpB* mutants (Figure 7B). This indicates that KhpA and KhpB might regulate bacterial stress responses in the stationary growth phase.

**Figure 7.**
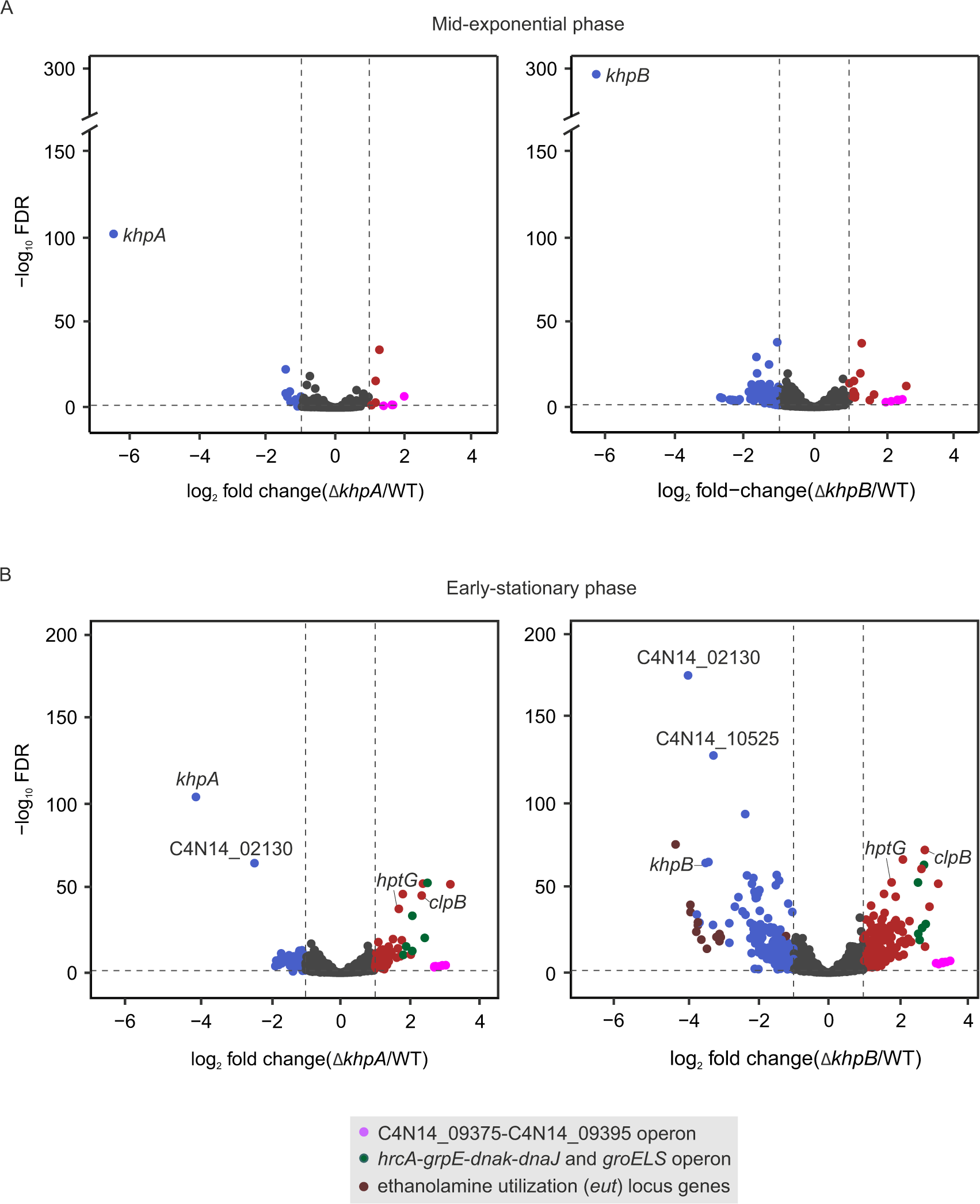
KhpA and KhpB regulate gene expression. (**A-B**) Volcano plots showing differential expression of all genes plotted against the corresponding FDR value in Δ*khpA* and Δ*khpB* strains in the mid-exponential growth phase (A) or early-stationary phase (B). Genes with a log_2_ fold change ≥1 and FDR ≤ 0.05 are considered upregulated and highlighted in red, those with a log_2_ fold-chang ≥1 and FDR ≤ 0.05 are downregulated and shown in blue. Specific transcripts and operons are labelled.

In line with the suggestion that KhpA and KhpB function as a heterodimer, we found overlapping sets of differentially expressed genes in both deletion strains during both growth phases. In the mid-exponential phase, all the differentially expressed transcripts in Δ*khpA* also show differential expression in Δ*khpB.* This includes genes from the operon C4N14_09080- C4N14_09105, which encodes the putative phosphotransferase system (PTS) and is among the strongest downregulated genes in Δ*khpA* and Δ*khpB* during this growth phase (Supplementary Fig. 6A, Supplementary Table S3). In the early stationary phase, 104 among the 117 differentially expressed transcripts in Δ*khpA* are also regulated by KhpB (Supplementary Fig. 6B, Supplementary Table S3). Interestingly, we noticed that 13 genes from an ethanolamine utilization (*eut*) locus are significantly downregulated in both deletion strains in the stationary phase (see below). Overall, these data suggest that KhpA and KhpB influence gene expression in *F. nucleatum*.

### KhpA and KhpB affect cell length

To characterize the cellular role of KhpA and KhpB, we first investigated their effect on cellular morphology, since KhpA was initially described as a protein that affects cell elongation in *S. pneumoniae* (34). Subsequent studies have confirmed this function in different species. In both *S. pneumoniae* and *Lactobacillus plantarum*, deletion or knockdown of *khpA* and *khpB* leads to decreased cell size (49–51). To examine if this function is conserved in *F. nucleatum*, we characterized cellular morphology of all deletion strains. Specifically, we stained the nucleus and membrane of cells from mid-exponential phase and measured the individual cell length (Figure 8A). We observed that the lack of either KhpA, KhpB or both leads to shortened cell length compared to WT, with the Δ*khpA*Δ*khpB* double mutant displaying the strongest effect (Figure 8B). To restore the expression of these proteins, we used the same strategy that we used to generate the Δ*khpB^khpB^*^-*3xFLAG*^ strain and generated an Δ*khpA^khpA^*^-*3xFLAG*^ strain. Complementation of both proteins in this manner rescued this effect, confirming that it is dependent on the function of both proteins and not caused by polar effects of the gene deletions (Figure 8B). These data show that the role of KhpA and KhpB in regulating cell length is conserved in *F. nucleatum*, despite its large phylogenetic distance to common model bacteria.

**Figure 8.**
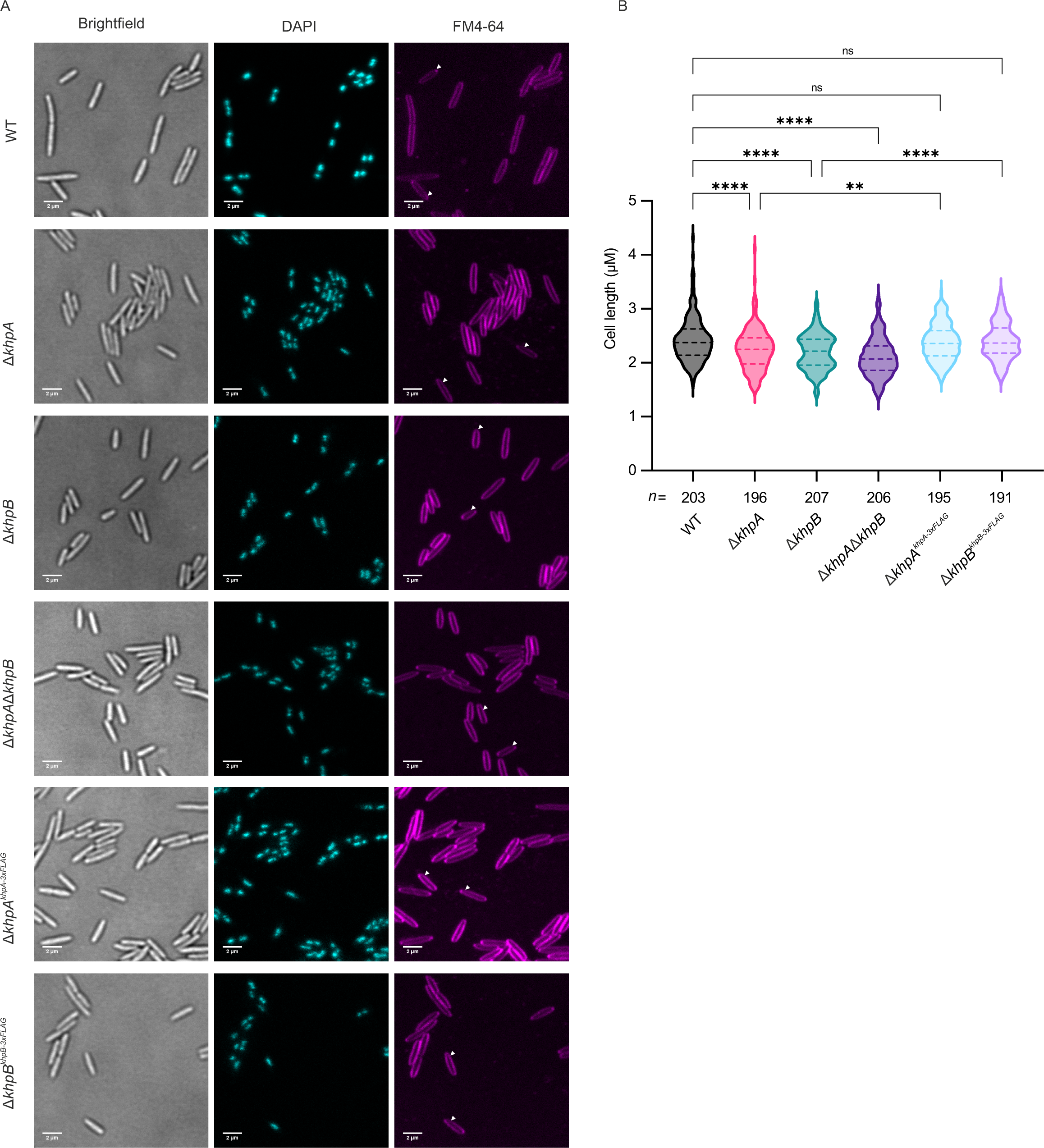
KhpA and KhpB affect cell length. (**A**) Microscopy of WT, deletion strains (Δ*khpA*, Δ*khpB*, Δ*khpA*Δ*khpB*) and complemented strains (Δ*khpA^khpA-3xFLAG^,* Δ*khpB^khpB-3xFLAG^*) in mid-exponential phase stained with DAPI and the membrane dye FM4-64. Shown are representative fields of vision for each strain from three biological replicates. For quantification, only cells with a single nucleoid based on DAPI staining were considered (indicated by white triangle). Scale bars =2 µm. (**B**) Violin plots of cell length measured from the microscopy images. The number of individual cells that were measured for each strain are indicated. The central dotted line of violin plot is the median value, upper and lower dotted lines are the interquartile range (IQR). P values were obtained relative to WT or Δ*khpA*, Δ*khpB* by one-way ANOVA analysis (GraphPad Prism). ns, not significant, \*\**P*<0.01, \*\*\*\**P*<0.0001.

### Loss of KhpA and KhpB impacts growth under nutrient limitation

To further characterize the physiological roles of KhpA and KhpB, we investigated their expression pattern during *F. nucleatum* growth in rich medium. We found that both proteins are expressed throughout the different growth phases and that they are upregulated upon entry into the mid-exponential growth phase (Figure 9A and 9B).

**Figure 9.**
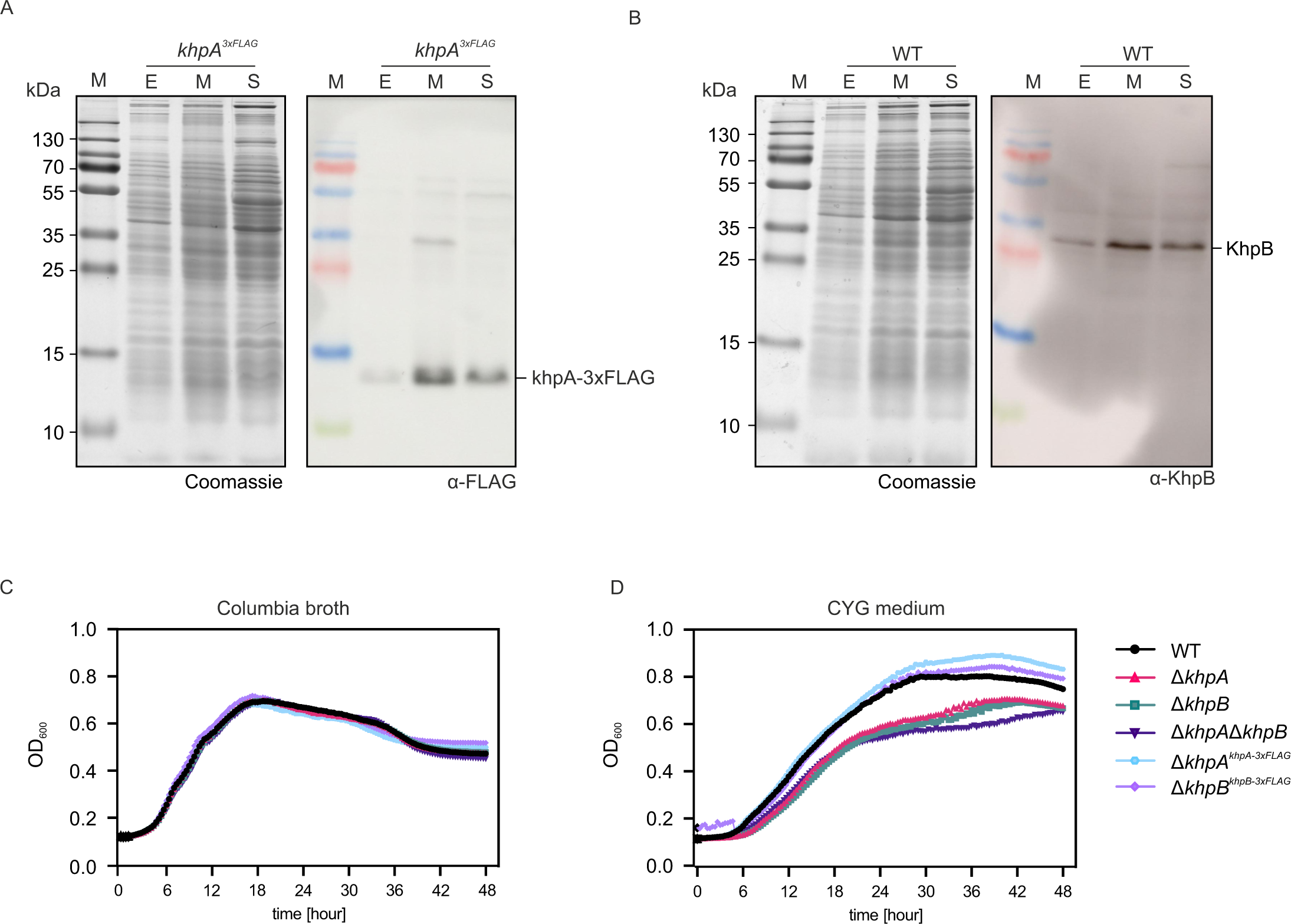
Loss of KhpA and KhpB impacts growth under nutrient limitation. (**A-B**) Western blot analysis of KhpA (A) and KhpB (B) expression during different growth phases. KhpB was detected with a KhpB specific antibody in the WT strain. KhpA was detected with a FLAG specific antibody in the *khpA^3xFLAG^* strain. Coomassie staining is used as loading control. E: early-exponential phase (OD_600_= 0.1), M: mid-exponential phase (OD_600_= 0.5), S: early-stationary phase (OD_600_= 1). (**C-D**) Growth curves for WT, deletion (Δ*khpA,* Δ*khpB,* Δ*khpA*Δ*khpB*) and complementedstrains (Δ*khpA^khpA-3xFLAG^,* Δ*khpB^khpB-3xFLAG^*) in Columbia broth (B) or CYG medium (C). Mean values of three biological replicates are plotted.

A growth comparison between WT, the individual deletion strains (Δ*khpA* or Δ*khpB*) and the double deletion strain (Δ*khpA*Δ*khpB*) in rich medium (Columbia broth) showed similar growth behavior (Figure 9C). In contrast, growth in CYG medium, a defined medium with only basal nutrients (see ‘Methods and Materials’), was negatively impacted in all deletion strains (Δ*khpA*, Δ*khpB* and Δ*khpA*Δ*khpB*), as indicated by a delay in growth and a reduced final optical density (Figure 9D). Importantly, restoring expression of KhpA (Δ*khpA^khpA-3xFLAG^*) or KhpB (Δ*khpB^khpB-3xFLAG^*) rescued this growth defect (Figure 9D). Overall, these observations suggest that KhpA and KhpB are not required under optimal growth conditions, but become important when the bacteria are deprived of nutrients. Taken together, KhpA and KhpB are likely involved in the response to nutrient stress in *F. nucleatum*.

### KhpB is involved in the regulation of ethanolamine utilization

Our RNA-seq analysis revealed the downregulation of 13 genes of the *eut* locus in the early stationary phase upon deletion of *khpA* or *khpB*, with a stronger effect upon *khpB* deletion (Figure 10A). The *eut* locus encodes enzymatic, structural, and regulatory components required for ethanolamine utilization (52). This pathway is of particular interest since ethanolamine is abundant in the gastrointestinal tract, represents an important nitrogen and carbon source (53) and might therefore play a role during the colonization of gastrointestinal cancer tissue by *F. nucleatum*. The regulated genes are located downstream of the putative *eutW* homolog of the *eut* locus, which functions as the sensory kinase of a two-component system with its upstream regulator *eutV* (Figure 10A). The EutVW system is activated upon ethanolamine binding and disrupts terminators present in the *eut* locus, allowing transcriptional read-through and expression of downstream genes as studied in other species (54–56).

**Figure 10.**
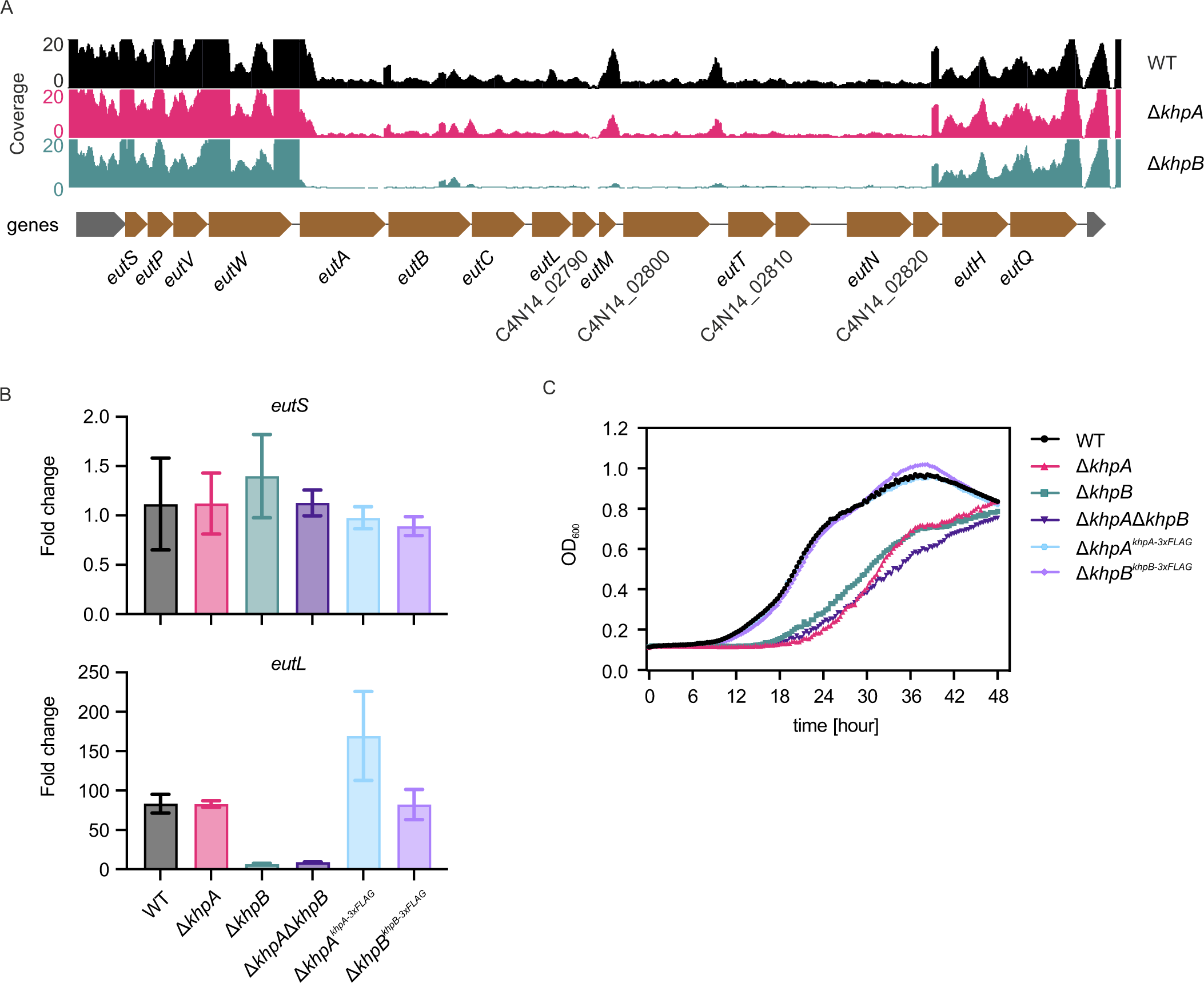
KhpB is involved in the regulation of ethanolamine utilization. (**A**) RNA-seq coverage on forward strand of the *eut* operon in WT, Δ*khpA* and Δ*khpB* (upper panel). Gene loci of *eut* operon (lower panel). (**B**) The *eutS* and *eutL* gene were evaluated by qRT-PCR in the indicated strains after transferring the cultures in ethanolamine-containing CYG medium for 2h. Fold change was normalized to the control, which is the RNA level in each indicated strain after growth 2h in CYG medium without ethanolamine, and plotted as the mean ± SD of three replicates. (**C**) Growth curves for WT, deletion (Δ*khpA,* Δ*khpB,* Δ*khpA*Δ*khpB*) and complemented strains (Δ*khpA^khpA-3xFLAG^,* Δ*khpB^khpB-3xFLAG^*) in CYG medium supplemented with 25 µM ethanolamine. Mean values of three replicates are plotted.

To validate the effect of *khpA or khpB* deletion on the expression of *eut* genes in the presence of ethanolamine, we transferred *F. nucleatum* from rich medium (Columbia broth) to CYG medium containing ethanolamine at the late-exponential phase and collected RNA samples after 2h. We then performed RT-qPCR analysis of *eutS* and *eutL*, two genes located upstream and downstream of *eutW*, respectively. While *eutS* levels are similar in all strains, *eutL* levels are >12 fold lower in the Δ*khpB* and >9 fold lower in Δ*khpA*Δ*khpB* background compared to WT (Figure 10B). Deletion of *khpA* did not have an effect on *eutL* levels. The relative RNA levels of *eutL* could be restored by chromosomal complementation of *khpB* or *khpA* (Figure 10B). These data indicate that KhpB is necessary for the transcription and/or stability of the *eutL* mRNA in *F. nucleatum*.

Next, we investigated the physiological relevance of the dysregulation of the *eut* genes in the deletion strains. To this end, we compared the growth of the WT and the different deletion strains (Δ*khpA*, Δ*khpB*, Δ*khpA*Δ*khpB)* in CYG medium supplemented with ethanolamine. We observed a pronounced growth delay (∼12 h) in all deletion strains compared to the WT (Figure 9C). While deletion of either *khpA* or *khpB* caused a similar growth delay, dysregulation of the *eut* locus in the Δ*khpA* strain was much weaker, an observation we currently do not yet understand. Nonetheless, complementation of either gene restored growth to WT levels (Figure 10C).

Overall, the results suggest that KhpB is required for a functional ethanolamine metabolism in *F. nucleatum*, indicating that the protein might be important during the colonization of secondary sites. Moreover, since this phenotype can be fully rescued by complementation of the *khpA* or *khpB* genes, it will provide an important departure point for setting up genetic screens to dissect the molecular architecture of the KhpA/B proteins and their potential interaction partners *in vivo*.

## DISCUSSION

This study describes an experimental search for sRNA-associated RBPs in *F. nucleatum*, a species of the poorly investigated bacterial phylum Fusobacteriota. Despite their poor genetic tractability, these bacteria are of particular interest for both their emerging medical importance and their phylogenetic vicinity to the last common ancestor of bacteria. The former motivates seeking a better understanding of fusobacterial gene regulation mechanisms, with the long-term goal to exploit such knowledge for the targeted elimination of *F. nucleatum* in secondary niches, such as colorectal or breast cancer tissue. The latter promises new insights into how RNA-mediated gene regulation through the activity of noncoding RNAs and RBPs evolved in prokaryotes. In this regard, our discovery of KhpA and KhpB as major sRNA-associated RBPs in *F. nucleatum* and their involvement in different biological processes sheds light on post-transcriptional regulation in the absence of the common bacterial RBPs CsrA, Hfq and ProQ.

## 6S RNA-RNAP complex formation is conserved in F. nucleatum

6S RNA-RNAP interactions have been detected in many bacteria (40) and our data show that *F. nucleatum* 6S RNA also interacts with the RNAP holoenzyme. Thus, we propose that the 6S-RNAP complex is likely ubiquitous in bacteria and can be generally used as a positive control in RNA interactome studies. Together with the previous detection of pRNAs in *F. nucleatum* (7), the 6S RNA-RNAP interaction also suggests that the formation of 6S RNA-RNAP complex, and by inference, RNA-mediated control of the transcription machinery, was established very early during the evolution of bacteria.

### New RBP candidates in fusobacteria

Our global screen identified a wealth of RBP candidates in *F. nucleatum*. Among these proteins, two stood out, because they interacted with many of the sRNA baits. One was the 3’-5’ exonuclease PNPase, which has previously been shown to be involved in regulating sRNA stability and function (57–59). Since the sRNA baits that enriched PNPase also pulled-down KhpA/B, it is tempting to speculate that PNPase is involved in regulating the stability of KhpA/B-interacting sRNAs in *F. nucleatum*. The other was the 30S ribosomal subunit protein RpsK. Due to its genomic locus and expression levels, the annotation of RpsK as a ribosomal protein is likely correct and further investigation will dissect why RpsK exhibits sRNAs-binding activity or if it associates with other sRNA-binding proteins in *F. nucleatum*.

We also recovered proteins involved in metabolic processes, consistent with prior RBP discovery studies that suggested that many metabolic enzymes moonlight as RBPs (45,60). For example, we find that the acyl carrier protein AcpP, which has a crucial role in bacterial fatty acid synthesis, was recovered by the majority of *F. nucleatum* sRNA baits. Curiously, the AcpP protein was also listed as a strong RBP candidate in *E. coli* based on analysis of the RNA bound proteome using global methods, such as TRAPP (total RNA-associated protein purification) (61) and RIC (RNA-interactome capture) (60). These observations suggest that AcpP may serve as an RBP in different bacteria, in addition to its core function in fatty acid biosynthesis.

Moreover, we identified proteins that had previously been reported as regulatory RBPs in other bacterial species. In addition to KhpA and KhpB, which will be discussed in the next section, the putative cold shock protein C4N14_09590 interacts with 14 sRNAs. Cold shock proteins (CSPs) are a family of small proteins that harbor a conserved RNA-binding cold shock domain (CSD) (62). They promote bacterial survival in lower temperatures and various other stress conditions by modulating transcription, translation and mRNA stability of target genes. CSPs have also been shown to be important for pathogenicity and metabolism (63–65). Therefore, it would be interesting to test if C4N14_09590 is indeed a functional CSP and if it might help *F. nucleatum* to adapt to different sites in human body.

In contrast to the broad sRNA interactors, the septation protein SpoVG binds only two sRNAs, FunR12 and FunR23, suggesting specificity. SpoVG is a widely conserved RBP in Gram-positive bacteria. It has recently been shown to be a post-transcriptional gene regulator similar to CsrA in Gram-negative bacteria and to affect carbon metabolism, biofilm formation and virulence in *Listeria monocytogenes* (66,67). It would therefore be interesting to determine the functional relevance of the SpoVG-sRNA interaction in *F. nucleatum*.

### KphA/B are major sRNA-binding proteins

Our global screen of sRNA-interactors identified KhpA and KhpB as associated with most of the sRNA baits in our study. Moreover, our RIP-seq analysis revealed that KhpA and KhpB bind multiple sRNAs and act together as global RBPs in *F. nucleatum*. However, the general principles that govern RNA binding of KhpA and KhpB are currently unclear. The KH domain in eukaryotes typically recognizes four nucleotides of single-stranded RNA, in which adenine bases hydrogen bond to the protein backbone (68). Given that KhpB forms a dimer with KhpA and possesses a second sequence-specific single-stranded-RNA-binding domain (R3H) (69), it was proposed that KhpA and KhpB interact with larger RNA regions than single KH domains. It remains unclear if KhpA and KhpB have preferential binding regions within a transcript, such as the 5’-UTR, the CDS or the 3’-UTR. Working out their binding preferences will be important to understand which regulatory networks they are involved in. For example, Hfq tends to bind both 5’ and 3’ UTRs of mRNAs to influence mRNA translation or stability (18,37,70), while ProQ shows preference to 3’ UTRs, where it counteracts exoribonuclease-dependent decay (25). Further studies applying global binding sites mapping approaches, such as CLIP-seq or CRAC (18,71) will be invaluable to better understand the main principles governing KhpB-RNA interactions and to help uncover the roles of KhpB in post-transcriptional gene regulation.

We found that KhpB affects the stability of several sRNAs. Similarly, *C. difficile* KhpB also influences the stability of several sRNAs both positively and negatively (35). Interestingly, KhpA and KhpB bind to the RNA degradosome in *Mycobacterium tuberculosis* (72), further suggesting a role in RNA turnover. As to next steps, global analysis of transcript decay in rifampicin run-on experiments should provide more evidence for whether KhpA and KhpB act as general stability factors *in F. nucleatum*, alike Hfq and ProQ (25,33,73). Interestingly, KhpA and KhpB both display a conserved genomic synteny with genes involved in RNA processing, such as *trmD* or *rnpA* (47). In *E. faecalis*, KhpB exhibits a potential role in processing tRNAs and sRNAs (36). These observations indicate that KhpB might play a role of RNA processing in different bacteria.

### Physiological roles of KphA/B

The physiological roles of KhpA and KhpB are beginning to be explored in several Gram-positive bacteria belonging to the phylum Firmicutes. In *C. difficile, S. pneumoniae* and *L. plantarum* both proteins are regulating cell elongation (34,35,49,51,74). *F. nucleatum khpA* or *khpB* deletion strains also display decreased cell length, strongly indicating a conserved role of KhpA and KhpB in controlling cell elongation across different phyla. While the underlying mechanism is not understood, it was speculated that KhpA and KhpB post-transcriptionally regulate expression of the cell division protein FtsA via the 5’-UTR of its mRNA (34). Interestingly, we observed that the transcripts of the cell division proteins FtsL and FtsZ immunoprecipitated with *F. nucleatum* KhpA and KhpB. However, we did not observe changes in the RNA levels of the *ftsL* and *ftsZ* mRNAs in *khpA* or *khpB* deletion strains. Nevertheless, it is possible that KhpA and KhpB affect the expression of FtsL and FtsZ via direct effects on translation.

Our data also demonstrate that KhpB is important for the regulation of ethanolamine utilization. This is indicated by the growth defect of *F. nucleatum khpA* or *khpB* deletion strains when ethanolamine is the main carbon source. Importantly, our transcriptomic data show that KhpB is required for the expression of many *eut* genes in the stationary growth phase. This suggests that KhpB might directly regulate the transcription or stability of these genes. By contrast, RIP-seq did not show a significant enrichment of these *eut* transcripts. Therefore, the precise regulatory mechanism requires further investigation. Nevertheless, by identifying a role of KhpB in regulating the utilization of the abundant intestinal metabolite ethanolamine in *F. nucleatum*, we have taken a first step to investigate the physiological function of this RBP as a gene regulator, which may play an important role for *F. nucleatum* colonization of secondary sites in human body.

### A robust phenotype for the molecular dissection of KhpA and KhpB residues

Phenotype-based forward genetic screens are a powerful strategy for determining important amino acid residues of a protein of interest (75). For instance, recent work took advantage of the observation that ProQ deficiency alleviates growth suppression of *Salmonella* with succinate as the sole carbon source to set up a mutagenesis screen. By coupling mutational scanning with loss-of-function selection, residues that are crucial for ProQ function and RNA binding have been determined (76). The fact that the *F. nucleatum khpB* deletion strain exhibits a strong growth defect in medium supplemented with ethanolamine can be exploited in a similar fashion to identify amino acid residues that are important for its function and to determine the motifs or residues that participate in binding to RNA ligands and in protein dimerization with KhpA.

### Are KH proteins a new family of sRNA chaperones in bacteria?

We have identified KhpA/B as major RBPs in *F. nucleatum* that bind to sRNAs and affect their stability. However, if KhpA/B have a similar role to Hfq in facilitating sRNAs base-pairing to target mRNAs remains unclear. Our study provides evidence for a direct interaction between KhpA/B and the sRNA FoxI, which is known to base-pair with target mRNAs to repress several envelope proteins in *F. nucleatum* (7). However, our preliminary data indicate that KhpB is not required for the FoxI-mediated repression of one of its targets. A global investigation of the KhpA/B-mediated RNA-RNA interactome by methods such as RIL-seq (RNA interaction by ligation and sequencing) (77) will determine if KhpA/B aid sRNA ligands base pair with their targets and serve as sRNA chaperones in the early-branching species *F. nucleatum*.

## DATA AVAILABILITY

The sequencing data have been deposited in the Gene Expression Omnibus (GEO) database https://www.ncbi.nlm.nih.gov/geo/query/acc.cgi?acc=GSE246396, (accession no. GSE246396). The mass spectrometry proteomics data have been deposited to the ProteomeXchange Consortium via the PRIDE partner repository with the dataset identifier PXD046416.

## AUTHOR CONTRIBUTIONS

Y.Z. performed most of the experiments. F.P. performed bioinformatic data analysis. V.C. conducted Confocal Microscopy assay. Y.Z, F.P and J.V. designed the research. Y.Z and J.V wrote the manuscript with input from F.P..

## ACKNOWLEDGEMENTS

We thank Anke Sparmann for comments on and editing of the manuscript; Kai Papenfort and Franziska Faber for comments on the manuscript; Esther Hauschild for collecting Confocal Microscopy images; Anna Zweyer for technical assist; Lars Schönemann for protein purification; Thorsten Bischler for RIP-seq data analysis; Stephanie Lamer for mass spectrometry analysis; The China Scholarship Council for awarding Y.Z. with a scholarship under the State Scholarship Fund; The Vogel Stiftung Dr. Eckernkamp for supporting F.P. and V.C. with a Dr. Eckernkamp Fellowship. The work was funded by a Gottfried-Wilhelm-Leibniz award to J.V. (DFG Vo875-18).

---

**Supplementary Figure S1.**
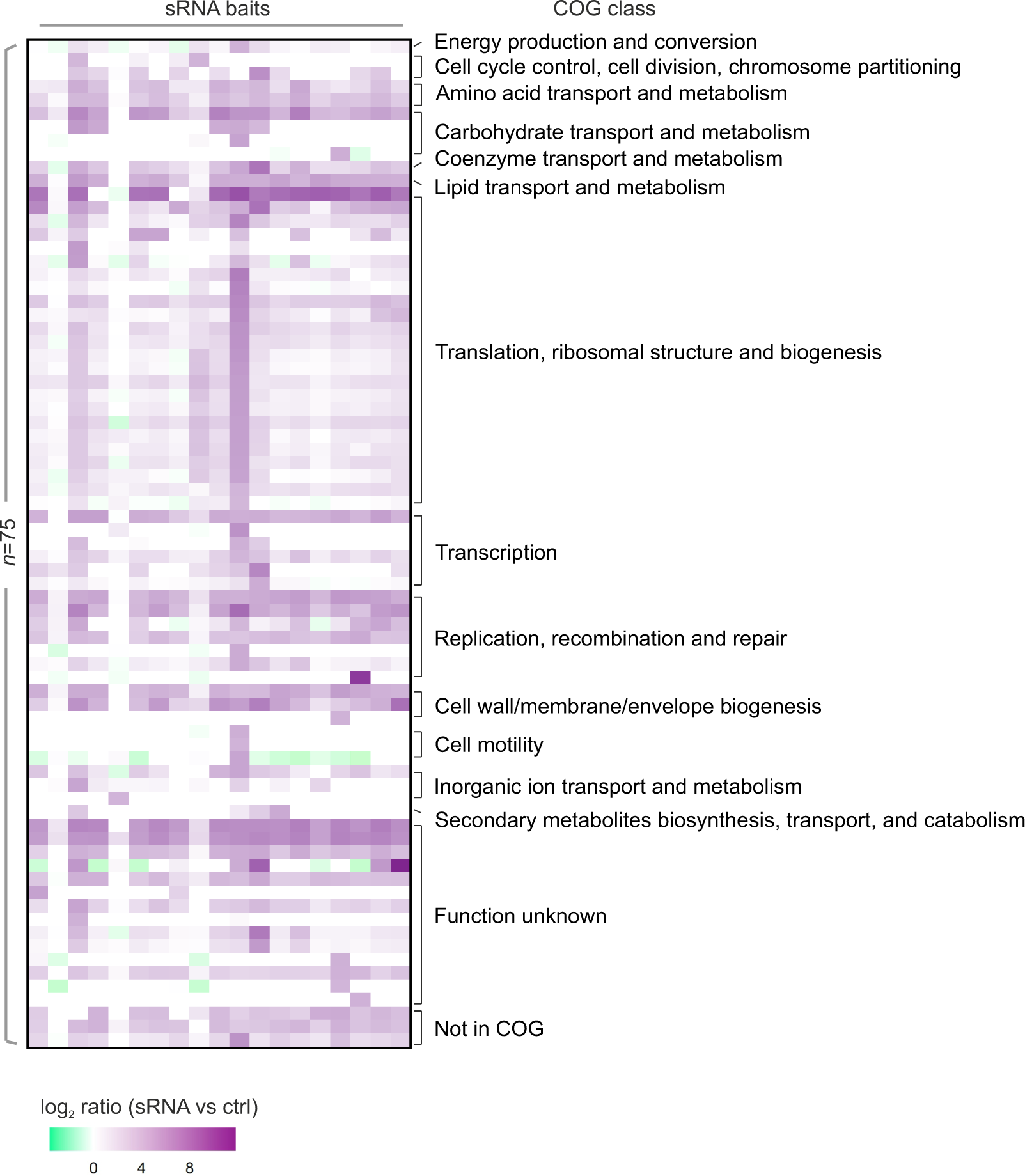
Significantly enriched proteins in sRNA affinity purification. Heat map representing log_2_ ratio of each significantly enriched protein (log_2_ ratio≥ 4) across all sRNA baits (n=75). The COG category of each protein is indicated.

**Supplementary Figure S2.**
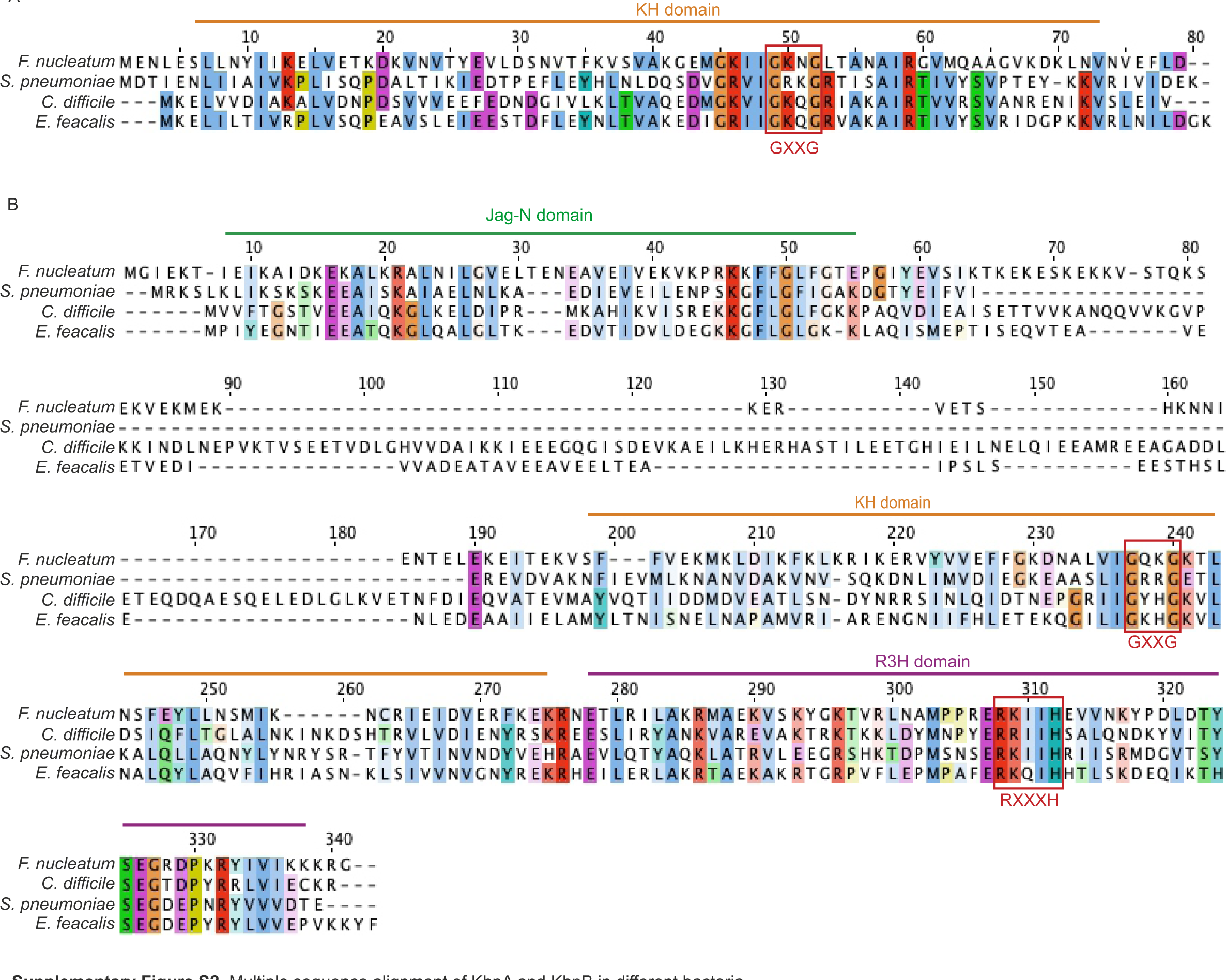
Multiple sequence alignment of KhpA and KhpB in different bacteria. (**A-B**) Sequences of the KhpA (A) and KhpB (B) from *F. nucleatum* and three other bacteria as indicated were subjected to multiple sequence alignment using Clustal Omega (Madeira *et al*., 2019) and were visualized using Jalview (Waterhouse *et al*., 2009). Residues with ≥ 50% identity are highlighted with colors. Domains and RNA-binding motifs are labeled.

**Supplementary Figure S3.**
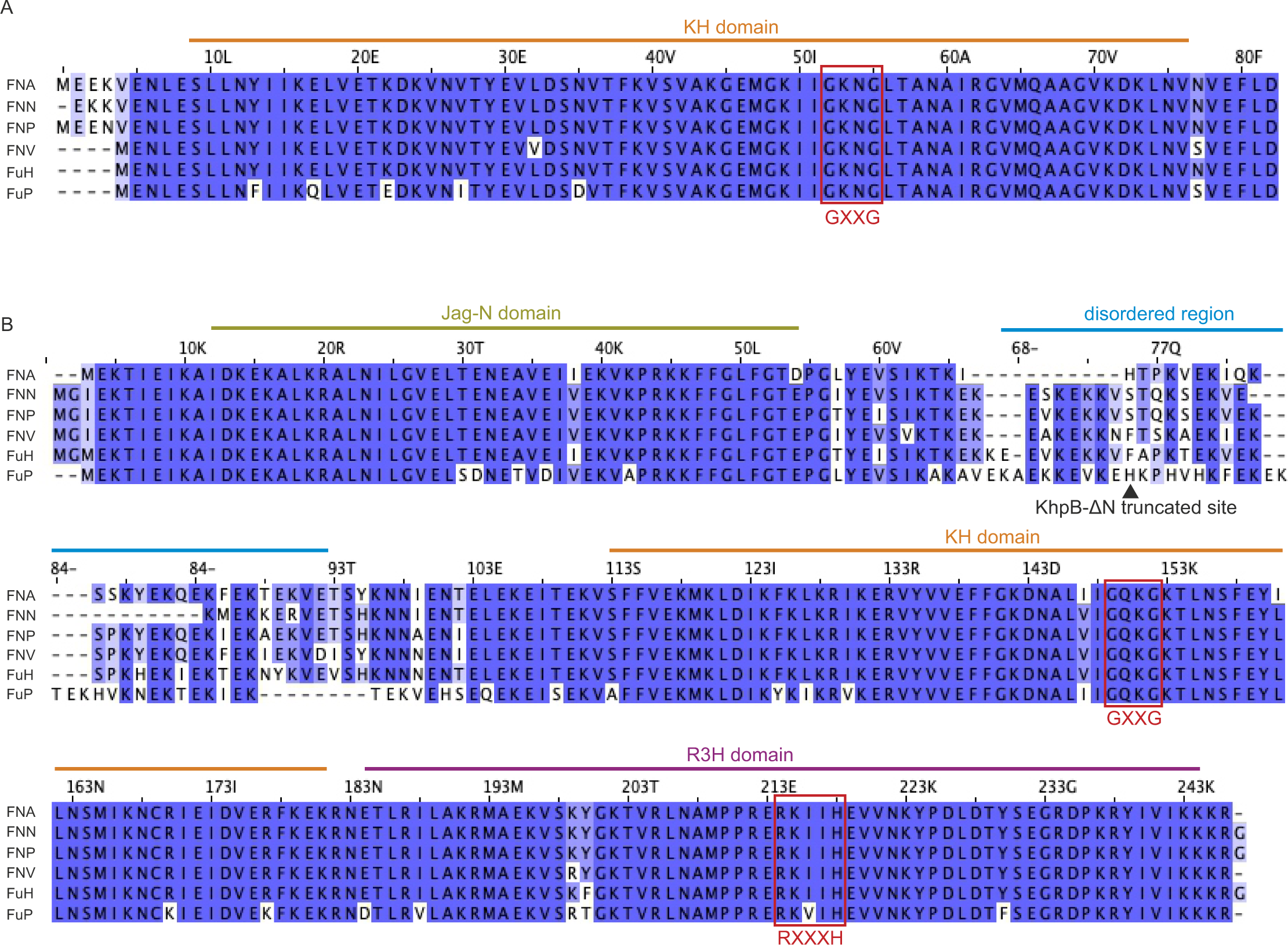
Multiple sequence alignment of KhpA and KhpB in *F. nucleatum* and related strains. (**A-B**) Sequences of the KhpA (A) and KhpB (B) from *F. nucleatum* and two related strains were subjected to multiple sequence alignment using Clustal Omega and were visualized using Jalview. Residues with ≥50% identity are highlighted in a blue gradient. FNN, *F. nucleatum subsp. nucleatum*; FNA, *F. nucleatum subsp. animalis*; FNP, *F. nucleatum subsp. polymorphum*; FNV, *F. nucleatum subsp. vincentii*; FuH, *F. hwasookii*; FuP, *F. periodonticum*. Domains and RNA-binding motifs are labeled. The truncation site that was used to generate purified KhpB-ΔN protein is indicated.

**Supplementary Figure S4.**
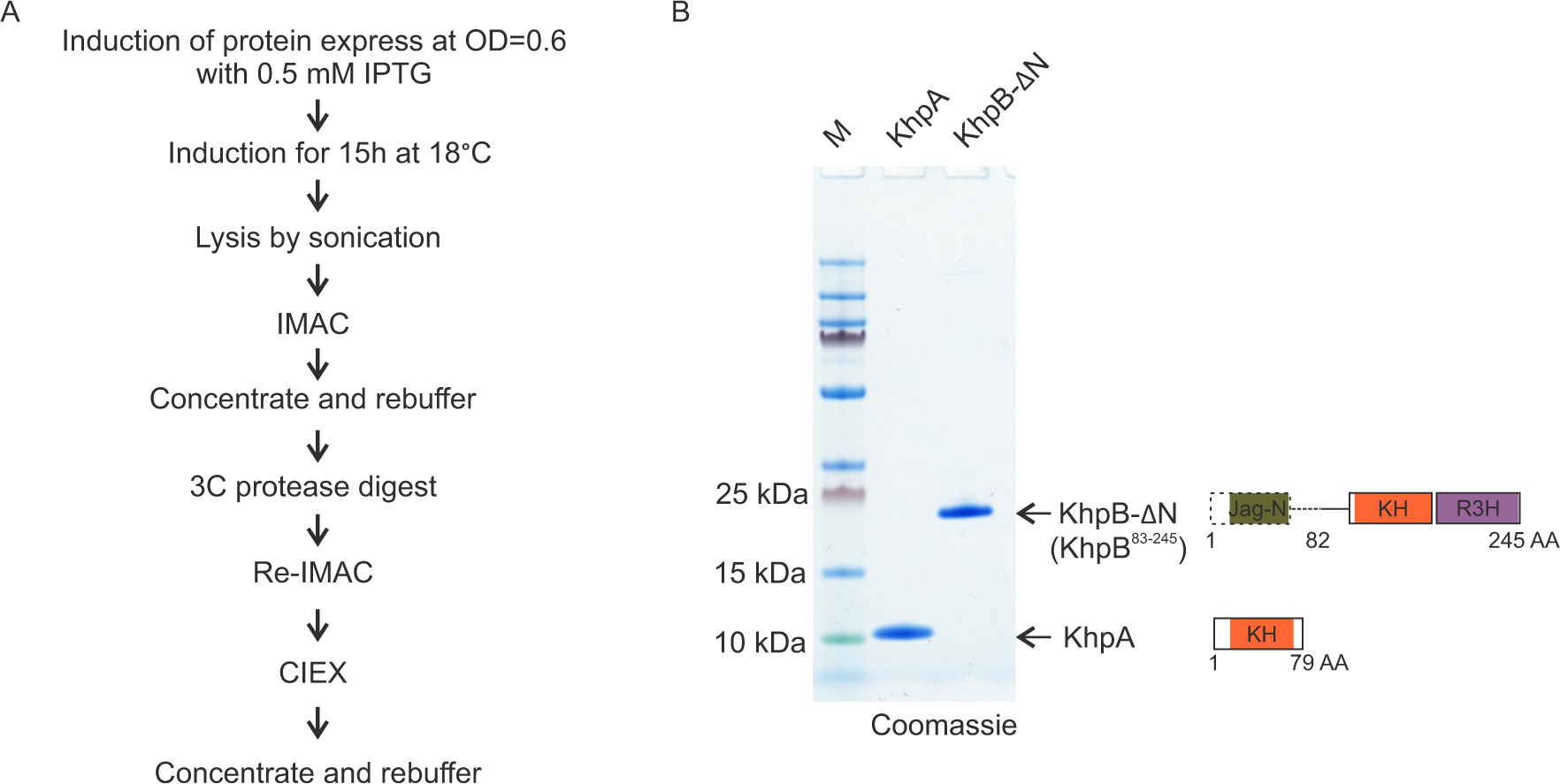
KhpA and KhpB-ΔN protein purification. (**A-B**) Purification scheme used to generate recombinant KhpA and KhpB-ΔN in *E. coli* (A). Protein purity after tag cleavage was checked by loading 2 µg of each protein on an SDS-PAGE gel that was stained by Coomassie (B).

**Supplementary Figure S5.**
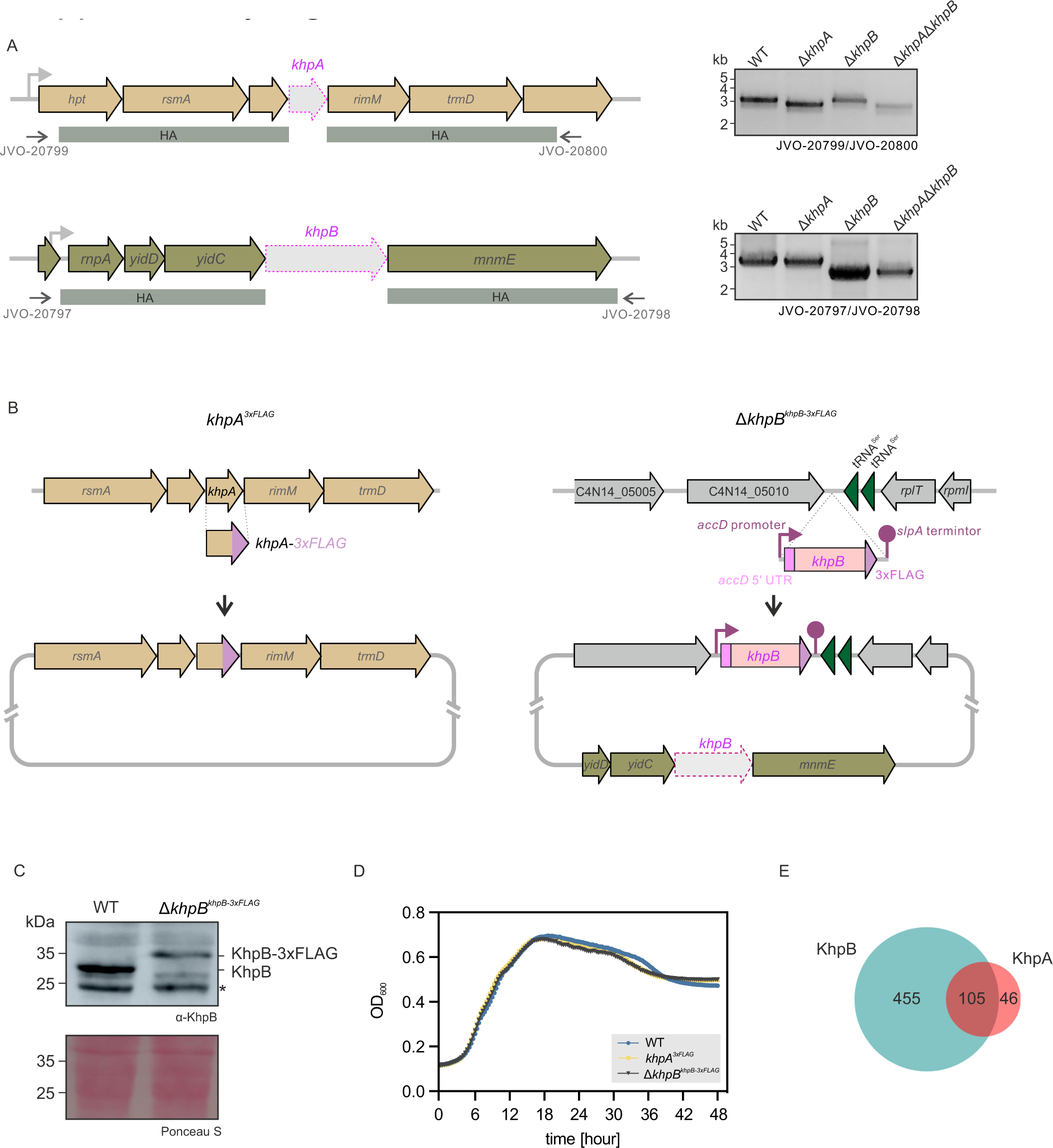
Deletion/tagging strains and RNA immunoprecipitation of KhpA and KhpB. (**A**) Schematic for *khpA* and *khpB* gene deletion and PCR verification. Upstream and downstream homologous recombination arms (HA) and primers used for PCR are indicated. Gel images of PCR verification of all deletion strains (Δ*khpA,* Δ*khpB,* Δ*khpA*Δ*khpB*) and WT using the indicated primers shown in the right panel. (**B**) Schematic for FLAG tagging *khpA* at its native locus (left) or *khpB* at an insulated complementation locus (right). (C)Western blot analysis of KhpB expression using a KhpB specific antibody in WT and Δ*khpB^khpB-3xFLAG^* strains. Ponceau S straining of the membrane after blotting is used to show equal loading. The asterisk indicates an unspecific band. (**D**) Growth curve for WT, *khpA^3xFLAG^* and Δ*khpB^khpB-3xFLAG^* strains in Columbia broth. Mean values of three replicates are plotted. (**E**) Venn diagram depicting the overlap of significantly enriched transcripts (log_2_ fold change ≥2) in KhpA and KhpB RIP-seq experiments.

**Supplementary Figure S6.**
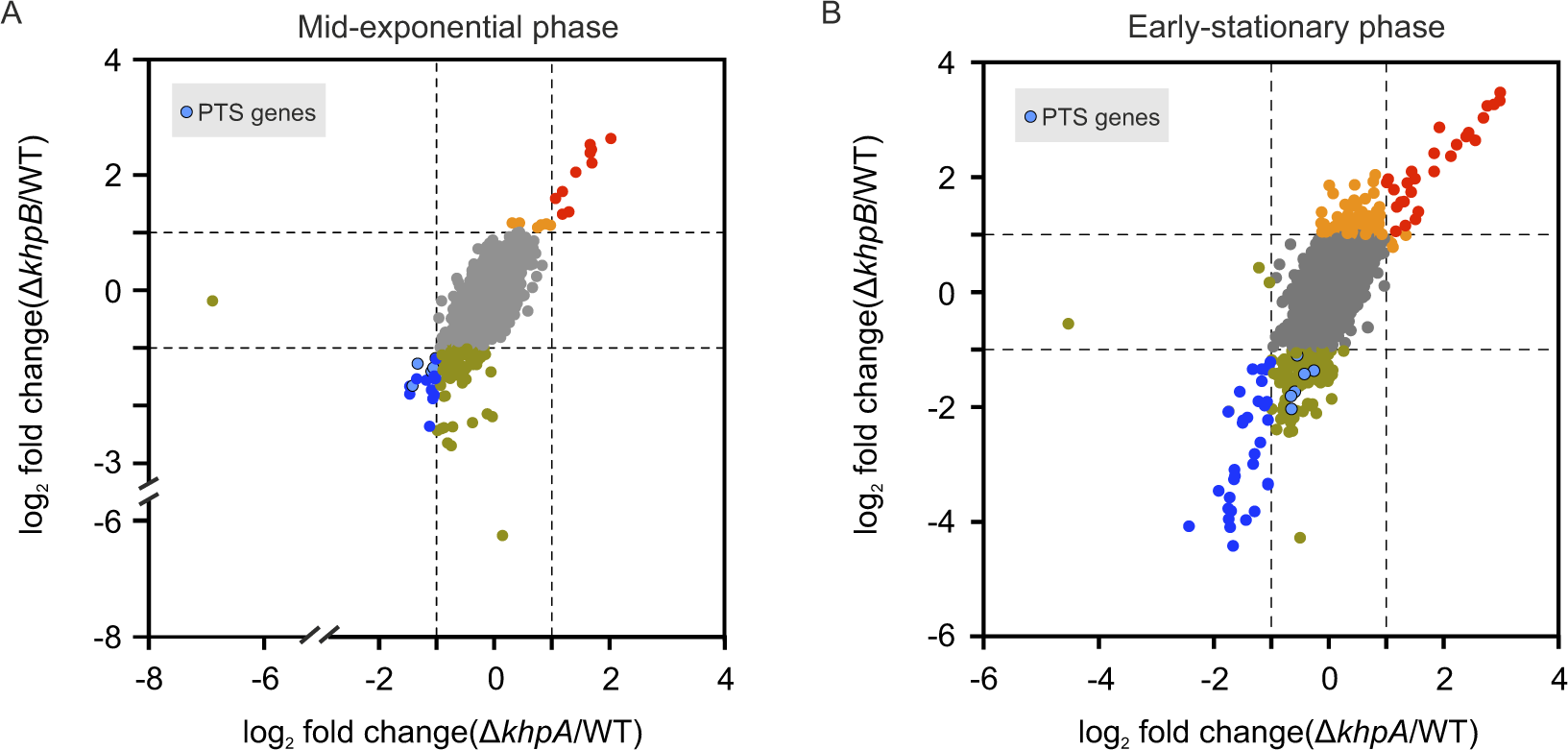
KhpA and KhpB regulate overlapping transcripts. (**A-B**)Scatter plots showing the log_2_ fold change for all transcripts in Δ*khpB* blotted against Δ*khpA* in mid-exponential phase (A) and early-stationary phase (B). PTS (phosphotransferase system) genes are highlighted.

## References

1. Brennan, C.A. and Garrett, W.S. (2019) Fusobacterium nucleatum - symbiont, opportunist and oncobacterium. Nat Rev Microbiol, 17, 156–166.

2. Han, Y.W. (2015) Fusobacterium nucleatum: a commensal-turned pathogen. Curr Opin Microbiol, 23, 141–147.

3. Wu, C., Chen, Y.W., Scheible, M., Chang, C., Wittchen, M., Lee, J.H., Luong, T.T., Tiner, B.L., Tauch, A., Das, A. et al. (2021) Genetic and molecular determinants of polymicrobial interactions in Fusobacterium nucleatum. Proc Natl Acad Sci U S A, 118.

4. Scheible, M., Nguyen, C.T., Luong, T.T., Lee, J.H., Chen, Y.W., Chang, C.Y., Wittchen, M., Camacho, M.I., Tiner, B.L., Wu, C.G. et al. (2022) The Fused Methionine Sulfoxide Reductase MsrAB Promotes Oxidative Stress Defense and Bacterial Virulence in Fusobacterium nucleatum. Mbio, 13.

5. Ponath, F., Zhu, Y., Cosi, V. and Vogel, J. (2022) Expanding the genetic toolkit helps dissect a global stress response in the early-branching species Fusobacterium nucleatum. Proc Natl Acad Sci U S A, 119, e2201460119.

6. Coleman, G.A., Davin, A.A., Mahendrarajah, T.A., Szantho, L.L., Spang, A., Hugenholtz, P., Szollosi, G.J. and Williams, T.A. (2021) A rooted phylogeny resolves early bacterial evolution. Science, 372.

7. Ponath, F., Tawk, C., Zhu, Y., Barquist, L., Faber, F. and Vogel, J. (2021) RNA landscape of the emerging cancer-associated microbe Fusobacterium nucleatum. Nat Microbiol, 6, 1007–1020.

8. Thompson, K.M., Rhodius, V.A. and Gottesman, S. (2007) SigmaE regulates and is regulated by a small RNA in Escherichia coli. J Bacteriol, 189, 4243–4256.

9. Papenfort, K., Pfeiffer, V., Mika, F., Lucchini, S., Hinton, J.C. and Vogel, J. (2006) SigmaE-dependent small RNAs of Salmonella respond to membrane stress by accelerating global omp mRNA decay. Mol Microbiol, 62, 1674–1688.

10. Melamed, S., Adams, P.P., Zhang, A., Zhang, H. and Storz, G. (2020) RNA-RNA Interactomes of ProQ and Hfq Reveal Overlapping and Competing Roles. Mol Cell, 77, 411–425 e417.

11. Sittka, A., Lucchini, S., Papenfort, K., Sharma, C.M., Rolle, K., Binnewies, T.T., Hinton, J.C. and Vogel, J. (2008) Deep sequencing analysis of small noncoding RNA and mRNA targets of the global post-transcriptional regulator, Hfq. PLoS Genet, 4, e1000163.

12. Chihara, K., Bischler, T., Barquist, L., Monzon, V.A., Noda, N., Vogel, J. and Tsuneda, S. (2019) Conditional Hfq Association with Small Noncoding RNAs in Pseudomonas aeruginosa Revealed through Comparative UV Cross-Linking Immunoprecipitation Followed by High-Throughput Sequencing. Msystems, 4.

13. Westermann, A.J., Venturini, E., Sellin, M.E., Forstner, K.U., Hardt, W.D. and Vogel, J. (2019) The Major RNA-Binding Protein ProQ Impacts Virulence Gene Expression in Salmonella enterica Serovar Typhimurium. mBio, 10.

14. Gulliver, E.L., Sy, B.M., Wong, J.L., Lucas, D.S.D., Powell, D.R., Harper, M., Tree, J.J. and Boyce, J.D. (2022) The Role and Targets of the RNA-Binding Protein ProQ in the Gram-Negative Bacterial Pathogen. Journal of Bacteriology, 204.

15. Ding, Y., Davis, B.M. and Waldor, M.K. (2004) Hfq is essential for Vibrio cholerae virulence and downregulates sigma expression. Mol Microbiol, 53, 345–354.

16. Holmqvist, E. and Vogel, J. (2018) RNA-binding proteins in bacteria. Nat Rev Microbiol, 16, 601–615.

17. Romeo, T., Gong, M., Liu, M.Y. and Brunzinkernagel, A.M. (1993) Identification and Molecular Characterization of Csra, a Pleiotropic Gene from Escherichia-Coli That Affects Glycogen Biosynthesis, Gluconeogenesis, Cell-Size, and Surface-Properties. Journal of Bacteriology, 175, 4744–4755.

18. Holmqvist, E., Wright, P.R., Li, L., Bischler, T., Barquist, L., Reinhardt, R., Backofen, R. and Vogel, J. (2016) Global RNA recognition patterns of post-transcriptional regulators Hfq and CsrA revealed by UV crosslinking in vivo. EMBO J, 35, 991–1011.

19. Dubey, A.K., Baker, C.S., Romeo, T. and Babitzke, P. (2005) RNA sequence and secondary structure participate in high-affinity CsrA-RNA interaction. RNA, 11, 1579–1587.

20. Weilbacher, T., Suzuki, K., Dubey, A.K., Wang, X., Gudapaty, S., Morozov, I., Baker, C.S., Georgellis, D., Babitzke, P. and Romeo, T. (2003) A novel sRNA component of the carbon storage regulatory system of Escherichia coli. Mol Microbiol, 48, 657–670.

21. Vogel, J. and Luisi, B.F. (2011) Hfq and its constellation of RNA. Nat Rev Microbiol, 9, 578–589.

22. Dimastrogiovanni, D., Frohlich, K.S., Bandyra, K.J., Bruce, H.A., Hohensee, S., Vogel, J. and Luisi, B.F. (2014) Recognition of the small regulatory RNA RydC by the bacterial Hfq protein. Elife, 3.

23. Moller, T., Franch, T., Hojrup, P., Keene, D.R., Bachinger, H.P., Brennan, R.G. and Valentin-Hansen, P. (2002) Hfq: a bacterial Sm-like protein that mediates RNA-RNA interaction. Molecular Cell, 9, 23–30.

24. Smirnov, A., Wang, C., Drewry, L.L. and Vogel, J. (2017) Molecular mechanism of mRNA repression in trans by a ProQ-dependent small RNA. EMBO J, 36, 1029–1045.

25. Holmqvist, E., Li, L., Bischler, T., Barquist, L. and Vogel, J. (2018) Global Maps of ProQ Binding In Vivo Reveal Target Recognition via RNA Structure and Stability Control at mRNA 3’ Ends. Mol Cell, 70, 971–982 e976.

26. Olejniczak, M. and Storz, G. (2017) ProQ/FinO-domain proteins: another ubiquitous family of RNA matchmakers? Mol Microbiol, 104, 905–915.

27. Gerovac, M., Vogel, J. and Smirnov, A. (2021) The World of Stable Ribonucleoproteins and Its Mapping With Grad-Seq and Related Approaches. Front Mol Biosci, 8, 661448.

28. Smith, T., Villanueva, E., Queiroz, R.M.L., Dawson, C.S., Elzek, M., Urdaneta, E.C., Willis, A.E., Beckmann, B.M., Krijgsveld, J. and Lilley, K.S. (2020) Organic phase separation opens up new opportunities to interrogate the RNA-binding proteome. Curr Opin Chem Biol, 54, 70–75.

29. Esteban-Serna, S., McCaughan, H. and Granneman, S. (2023) Advantages and limitations of UV cross-linking analysis of protein-RNA interactomes in microbes. Molecular Microbiology.

30. Gemmill, D., D’Souza, S., Meier-Stephenson, V. and Patel, T.R. (2020) Current approaches for RNA-labelling to identify RNA-binding proteins. Biochem Cell Biol, 98, 31–41.

31. Treiber, T., Treiber, N. and Meister, G. (2018) Identification of microRNA Precursor-Associated Proteins. Methods Mol Biol, 1823, 103–114.

32. 32. Hor, J., Garriss, G., Di Giorgio, S., Hack, L.M., Vanselow, J.T., Forstner, K.U., Schlosser, A., Henriques-Normark, B. and Vogel, J. (2020) Grad-seq in a Gram-positive bacterium reveals exonucleolytic sRNA activation in competence control. EMBO J, 39, e103852.

33. Smirnov, A., Forstner, K.U., Holmqvist, E., Otto, A., Gunster, R., Becher, D., Reinhardt, R. and Vogel, J. (2016) Grad-seq guides the discovery of ProQ as a major small RNA-binding protein. Proc Natl Acad Sci U S A, 113, 11591–11596.

34. Zheng, J.J., Perez, A.J., Tsui, H.T., Massidda, O. and Winkler, M.E. (2017) Absence of the KhpA and KhpB (JAG/EloR) RNA-binding proteins suppresses the requirement for PBP2b by overproduction of FtsA in Streptococcus pneumoniae D39. Mol Microbiol, 106, 793–814.

35. Lamm-Schmidt, V., Fuchs, M., Sulzer, J., Gerovac, M., Hör, J., Dersch, P., Vogel, J. and Faber, F. (2021) Grad-seq identifies KhpB as a global RNA-binding protein in Clostridioides difficile that regulates toxin production. microLife, 2.

36. Michaux, C., Gerovac, M., Hansen, E.E., Barquist, L. and Vogel, J. (2023) Grad-seq analysis of Enterococcus faecalis and Enterococcus faecium provides a global view of RNA and protein complexes in these two opportunistic pathogens. microLife, 4.

37. Chao, Y., Papenfort, K., Reinhardt, R., Sharma, C.M. and Vogel, J. (2012) An atlas of Hfq-bound transcripts reveals 3’ UTRs as a genomic reservoir of regulatory small RNAs. EMBO J, 31, 4005–4019.

38. Livak, K.J. and Schmittgen, T.D. (2001) Analysis of relative gene expression data using real-time quantitative PCR and the 2(-Delta Delta C(T)) Method. Methods, 25, 402–408.

39. Said, N., Rieder, R., Hurwitz, R., Deckert, J., Urlaub, H. and Vogel, J. (2009) In vivo expression and purification of aptamer-tagged small RNA regulators. Nucleic Acids Res, 37, e133.

40. Wassarman, K.M. (2018) 6S RNA, a Global Regulator of Transcription. Microbiol Spectr, 6.

41. Wassarman, K.M. and Storz, G. (2000) 6S RNA regulates E. coli RNA polymerase activity. Cell, 101, 613–623.

42. Wassarman, K.M. and Saecker, R.M. (2006) Synthesis-mediated release of a small RNA inhibitor of RNA polymerase. Science, 314, 1601–1603.

43. Ashburner, M., Ball, C.A., Blake, J.A., Botstein, D., Butler, H., Cherry, J.M., Davis, A.P., Dolinski, K., Dwight, S.S., Eppig, J.T. et al. (2000) Gene ontology: tool for the unification of biology. The Gene Ontology Consortium. Nat Genet, 25, 25–29.

44. Galperin, M.Y., Wolf, Y.I., Makarova, K.S., Vera Alvarez, R., Landsman, D. and Koonin, E.V. (2021) COG database update: focus on microbial diversity, model organisms, and widespread pathogens. Nucleic Acids Res, 49, D274–D281.

45. 45. Chu, L.C., Arede, P., Li, W., Urdaneta, E.C., Ivanova, I., McKellar, S.W., Wills, J.C., Frohlich, T., von Kriegsheim, A., Beckmann, B.M., et al. (2022) The RNA-bound proteome of MRSA reveals post-transcriptional roles for helix-turn-helix DNA-binding and Rossmann-fold proteins. Nat Commun, 13, 2883.

46. Hentze, M.W., Castello, A., Schwarzl, T. and Preiss, T. (2018) A brave new world of RNA-binding proteins. Nat Rev Mol Cell Biol, 19, 327–341.

47. Olejniczak, M., Jiang, X., Basczok, M.M. and Storz, G. (2022) KH domain proteins: Another family of bacterial RNA matchmakers? Mol Microbiol, 117, 10–19.

48. Nicastro, G., Taylor, I.A. and Ramos, A. (2015) KH-RNA interactions: back in the groove. Curr Opin Struct Biol, 30, 63–70.

49. Winther, A.R., Kjos, M., Stamsas, G.A., Havarstein, L.S. and Straume, D. (2019) Prevention of EloR/KhpA heterodimerization by introduction of site-specific amino acid substitutions renders the essential elongasome protein PBP2b redundant in Streptococcus pneumoniae. Sci Rep, 9, 3681.

50. Stamsas, G.A., Straume, D., Ruud Winther, A., Kjos, M., Frantzen, C.A. and Havarstein, L.S. (2017) Identification of EloR (Spr1851) as a regulator of cell elongation in Streptococcus pneumoniae. Mol Microbiol, 105, 954–967.

51. Myrbraten, I.S., Wiull, K., Salehian, Z., Havarstein, L.S., Straume, D., Mathiesen, G. and Kjos, M. (2019) CRISPR Interference for Rapid Knockdown of Essential Cell Cycle Genes in Lactobacillus plantarum. mSphere, 4.

52. Tsoy, O., Ravcheev, D. and Mushegian, A. (2009) Comparative genomics of ethanolamine utilization. J Bacteriol, 191, 7157–7164.

53. Garsin, D.A. (2010) Ethanolamine utilization in bacterial pathogens: roles and regulation. Nat Rev Microbiol, 8, 290–295.

54. Del Papa, M.F. and Perego, M. (2008) Ethanolamine activates a sensor histidine kinase regulating its utilization in Enterococcus faecalis. J Bacteriol, 190, 7147–7156.

55. Fox, K.A., Ramesh, A., Stearns, J.E., Bourgogne, A., Reyes-Jara, A., Winkler, W.C. and Garsin, D.A. (2009) Multiple posttranscriptional regulatory mechanisms partner to control ethanolamine utilization in Enterococcus faecalis. P Natl Acad Sci USA, 106, 4435–4440.

56. Baker, K.A. and Perego, M. (2011) Transcription antitermination by a phosphorylated response regulator and cobalamin-dependent termination at a B(1)(2) riboswitch contribute to ethanolamine utilization in Enterococcus faecalis. J Bacteriol, 193, 2575–2586.

57. 57. Cameron, T.A., Matz, L.M., Sinha, D. and De Lay, N.R. (2019) Polynucleotide phosphorylase promotes the stability and function of Hfq-binding sRNAs by degrading target mRNA-derived fragments. Nucleic Acids Res, 47, 8821–8837.

58. 58. Dendooven, T., Sinha, D., Roeselova, A., Cameron, T.A., De Lay, N.R., Luisi, B.F. and Bandyra, K.J. (2021) A cooperative PNPase-Hfq-RNA carrier complex facilitates bacterial riboregulation. Molecular Cell, 81, 2901-+.

59. Andrade, J.M., Pobre, V., Matos, A.M. and Arraiano, C.M. (2012) The crucial role of PNPase in the degradation of small RNAs that are not associated with Hfq. RNA, 18, 844–855.

60. Stenum, T.S., Kumar, A.D., Sandbaumhüter, F.A., Kjellin, J., Jerlström-Hultqvist, J., Andrén, P.E., Koskiniemi, S., Jansson, E.T. and Holmqvist, E. (2023) RNA interactome capture in Escherichia coli globally identifies RNA-binding proteins. Nucleic Acids Research, 51, 4572–4587.

61. Shchepachev, V., Bresson, S., Spanos, C., Petfalski, E., Fischer, L., Rappsilber, J. and Tollervey, D. (2019) Defining the RNA interactome by total RNA-associated protein purification. Mol Syst Biol, 15, e8689.

62. Horn, G., Hofweber, R., Kremer, W. and Kalbitzer, H.R. (2007) Structure and function of bacterial cold shock proteins. Cell Mol Life Sci, 64, 1457–1470.

63. Michaux, C., Holmqvist, E., Vasicek, E., Sharan, M., Barquist, L., Westermann, A.J., Gunn, J.S. and Vogel, J. (2017) RNA target profiles direct the discovery of virulence functions for the cold-shock proteins CspC and CspE. Proc Natl Acad Sci U S A, 114, 6824–6829.

64. 64. Muchaamba, F., von Ah, U., Stephan, R., Stevens, M.J.A. and Tasara, T. (2022) Deciphering the global roles of Cold shock proteins in Listeria monocytogenes nutrient metabolism and stress tolerance. Front Microbiol, 13, 1057754.

65. Jiang, W., Hou, Y. and Inouye, M. (1997) CspA, the major cold-shock protein of Escherichia coli, is an RNA chaperone. J Biol Chem, 272, 196–202.

66. Burke, T.P. and Portnoy, D.A. (2016) SpoVG Is a Conserved RNA-Binding Protein That Regulates Listeria monocytogenes Lysozyme Resistance, Virulence, and Swarming Motility. mBio, 7, e00240.

67. Shi, C.Z., Zheng, L.P., Lu, Z.X., Zhang, X.Y. and Bie, X.M. (2023) The global regulator SpoVG regulates Listeria monocytogenes biofilm formation. Microb Pathogenesis, 180.

68. Valverde, R., Edwards, L. and Regan, L. (2008) Structure and function of KH domains. FEBS J, 275, 2712–2726.

69. Grishin, N.V. (1998) The R3H motif: A domain that binds single-stranded nucleic acids. Trends Biochem Sci, 23, 329–330.

70. 70. Iosub, I.A., van Nues, R.W., McKellar, S.W., Nieken, K.J., Marchioretto, M., Sy, B., Tree, J.J., Viero, G. and Granneman, S. (2020) Hfq CLASH uncovers sRNA-target interaction networks linked to nutrient availability adaptation. Elife, 9.

71. Tree, J.J., Gerdes, K. and Tollervey, D. (2018) Transcriptome-Wide Analysis of Protein-RNA and RNA-RNA Interactions in Pathogenic Bacteria. Methods Enzymol, 612, 467–488.

72. Plocinski, P., Macios, M., Houghton, J., Niemiec, E., Plocinska, R., Brzostek, A., Slomka, M., Dziadek, J., Young, D. and Dziembowski, A. (2019) Proteomic and transcriptomic experiments reveal an essential role of RNA degradosome complexes in shaping the transcriptome of Mycobacterium tuberculosis. Nucleic Acids Res, 47, 5892–5905.

73. Bauriedl, S., Gerovac, M., Heidrich, N., Bischler, T., Barquist, L., Vogel, J. and Schoen, C. (2020) The minimal meningococcal ProQ protein has an intrinsic capacity for structure-based global RNA recognition. Nat Commun, 11, 2823.

74. Ulrych, A., Holeckova, N., Goldova, J., Doubravova, L., Benada, O., Kofronova, O., Halada, P. and Branny, P. (2016) Characterization of pneumococcal Ser/Thr protein phosphatase phpP mutant and identification of a novel PhpP substrate, putative RNA binding protein Jag. BMC Microbiol, 16, 247.

75. Fowler, D.M. and Fields, S. (2014) Deep mutational scanning: a new style of protein science. Nat Methods, 11, 801–807.

76. El Mouali, Y., Ponath, F., Scharrer, V., Wenner, N., Hinton, J.C.D. and Vogel, J. (2021) Scanning mutagenesis of RNA-binding protein ProQ reveals a quality control role for the Lon protease. Rna, 27, 1512–1527.

77. Melamed, S., Peer, A., Faigenbaum-Romm, R., Gatt, Y.E., Reiss, N., Bar, A., Altuvia, Y., Argaman, L. and Margalit, H. (2016) Global Mapping of Small RNA-Target Interactions in Bacteria. Mol Cell, 63, 884–897.

